# Climbing up and down binding landscapes: a high-throughput study of mutational effects in homologous protein-protein complexes

**DOI:** 10.1101/2020.10.14.338756

**Authors:** Michael Heyne, Jason Shirian, Itay Cohen, Yoav Peleg, Evette S. Radisky, Niv Papo, Julia M. Shifman

## Abstract

Each protein-protein interaction (PPI) has evolved to possess binding affinity that is compatible with its cellular function. As such, cognate enzyme/inhibitor interactions frequently exhibit very high binding affinities, while structurally similar non-cognate PPIs possess substantially weaker binding affinities. To understand how slight differences in sequence and structure could lead to drastic changes in PPI binding free energy (ΔΔ*G*_bind_), we study three homologous PPIs that span nine orders of magnitude in binding affinity and involve a serine protease interacting with an inhibitor BPTI. Using state-of-the-art methodology that combines protein randomization and affinity sorting coupled to next-generation sequencing and data normalization, we report quantitative binding landscapes consisting of ΔΔ*G*_bind_ values for the three PPIs, gleaned from tens of thousands of single and double mutations in the BPTI binding interface. We demonstrate that the three homologous PPIs possess drastically different binding landscapes and lie at different points in respect to the landscape maximum. Furthermore, the three PPIs demonstrate distinct patterns of coupling energies between two simultaneous mutations that depend not only on positions involved but also on the nature of the mutation. Interestingly, we find that in all three PPIs positive epistasis is frequently observed at hot-spot positions where mutations lead to loss of high affinity, while conversely negative epistasis is observed at cold-spot positions, where mutations lead to affinity enhancement. The new insights on PPI evolution revealed in this study will be invaluable in understanding evolution of other biological complexes and can greatly facilitate design of novel high-affinity protein inhibitors.

**Significance:** Protein-protein interactions (PPIs) have evolved to display binding affinities that can support their function. As such, cognate and non-cognate PPIs could be highly similar structurally but exhibit huge differences in binding affinities. To understand this phenomenon, we studied the effect of tens of thousands of single and double mutations on binding affinity of three homologous protease-inhibitor complexes. We show that binding landscapes of the three complexes are strikingly different and depend on the PPI evolutionary optimality. We observe different patterns of couplings between mutations for the three PPIs with negative and positive epistasis appearing most frequently at hot-spot and cold-spot positions, respectively. The evolutionary trends observed here are likely to be universal to all biological complexes in the cell.

## Introduction

Protein function is determined by the protein amino acid sequence, which has undergone millions of years of evolution while subjected to various selection pressures. Native proteins have evolved not only to perform their main function but also to satisfy a number of criteria such as solubility (1), low propensity for aggregation, stability, resistance to stress conditions (2), etc. As a result, proteins usually function below their maximum capacity (3). Multiple experiments on enzymes and binding domains proved that protein fitness could be enhanced by several orders of magnitude by applying an appropriate pressure and selecting the fittest protein sequences (4–7).

Fitness landscapes explore the effects of all possible mutations on the ability of proteins to perform their main function. Such landscapes reveal how far a particular protein is from its functional maximum, what fraction of mutations leads up and down the “fitness hill”, how large the mutational steps are and which residues are the most critical to protein function (8). Mapping of fitness landscapes is thus an attractive strategy for approaching various protein engineering projects with the goal to improve or modify protein function since the best mutations could be easily identified from the fitness landscape (9, 10). Development of new strategies for protein randomization and advances in Next Generation Sequencing (NGS) enabled several exciting studies that report fitness landscapes for a number of biological systems (1, 11–22). In these studies, the effects of mutations on enzyme catalysis, fluorescence, thermostability, and other functions have been reported, giving invaluable insights on how different biological functions have evolved.

One out of many important protein functions is binding between two or more protein partners. Binding is crucial in many cellular activities such as signal transduction, protein regulation, transcription/ translation and others. Mutations in protein-protein interactions (PPIs) frequently result in a change in free energy of binding (ΔΔG_bind_), sometimes weakening and sometimes stabilizing the interaction (23). A mutation resulting in substantial ΔΔG_bind_ in one PPI could translate into remodeling of the whole PPI network, frequently leading to dysregulation of signal transduction pathways and disease (24, 25). Therefore, understanding how mutations in PPIs affect their binding affinity is of great importance to both basic biology and to biomedical sciences, where inhibition or activation of a particular PPI might help to treat the related disease.

At the present moment, comprehensive binding landscapes have been mapped for only a handful of proteins(9, 26–30), while for most PPIs only a few ΔΔG_bind_ data points have been measured, most frequently corresponding to mutations to alanines (31–35). Comparison of the available sparse ΔΔG_bind_ data from different studies let us hypothesize that different classes of PPIs possess principally different binding landscapes and lie at different points relative to the binding landscape maximum, i. e. the amino acid sequence with the highest possible affinity. While in some PPIs, the majority of single mutations lead to large destabilization of the protein-protein complex (28, 36–38), in other PPIs frequent affinity-enhancing mutations are observed (27, 39). The magnitude of ΔΔG_bind_ due to mutation is likely to depend on the nature of the PPI under study as well as on the location of the mutation within the protein. It has been demonstrated that a few critical positions termed hot-spots of binding contribute the most significantly to the PPI binding energy with mutations at those positions usually leading to large drop in affinity (40–42). Coldspot positions, on the other hand, present multiple possibilities for PPI affinity improvement (43).

To investigate the relationship between PPI structure/function and the corresponding binding free energy, we set to compare comprehensive binding landscapes of three homologous PPIs that span nine orders in magnitude in binding affinity (K_D_). These complexes include Bovine Pancreatic Trypsin Inhibitor (BPTI) interacting with three serine proteases: Bovine trypsin (BT) (K_D_ = 10^-14^ M) (38), Bovine α-Chymotrypsin (ChT) (K_D_ = 10^-8^ M) (38) and Human Mesotrypsin (MT) (K_D_ = 10^-5^ M) (44). In spite of such large differences in K_D_s, the binding interfaces of the three complexes are highly similar, exhibiting nearly identical physicochemical properties (Figure 1).

**Figure 1:**
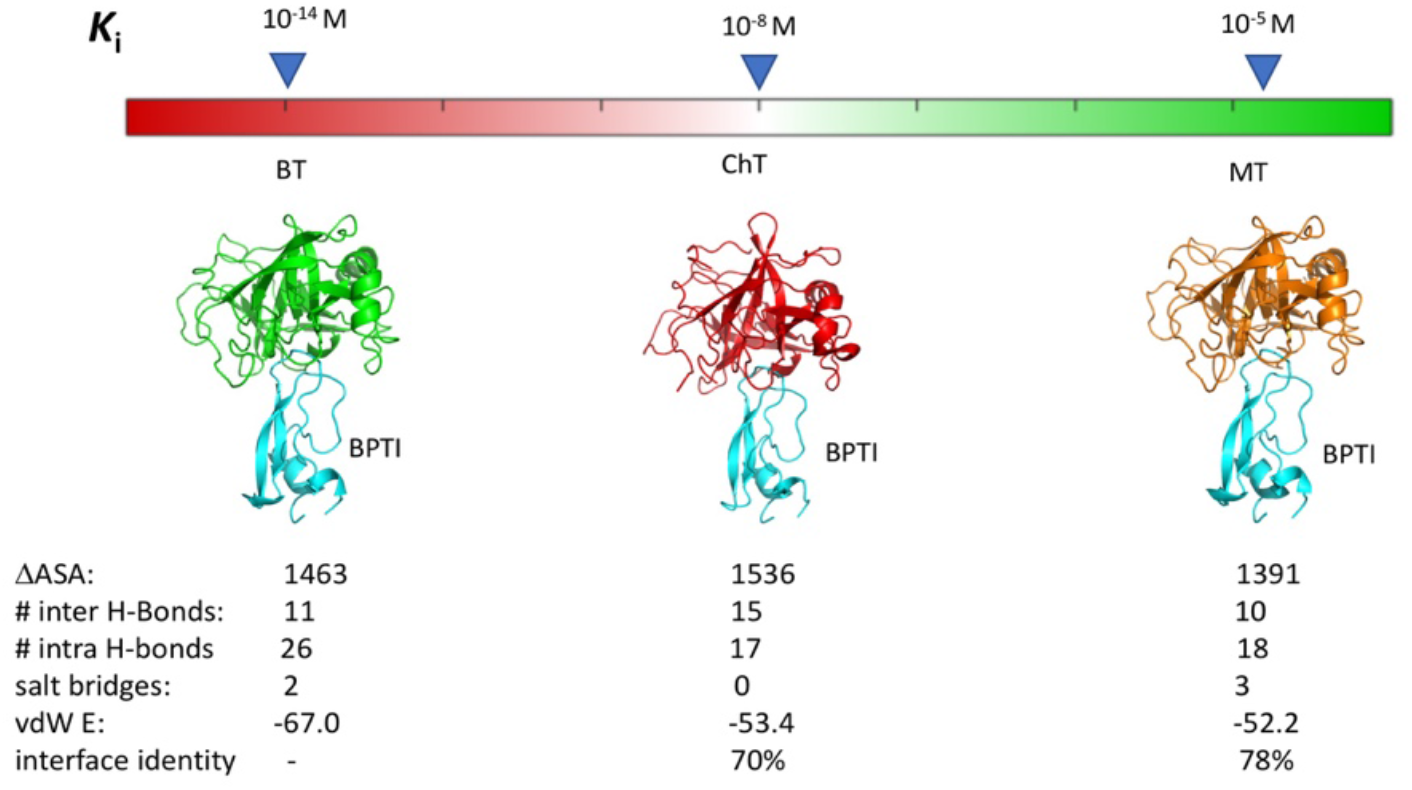
Comparison of K_D_s and structures for the 3 PPIs. Structures of the 3 complexes between BPTI and BT (PDB ID 3OTJ), ChT (PDB ID 1CBW) and MT (PDB ID 2R9P) are shown. Summarize are inhibition constants, change in solvent-accessible surface area upon complex formation (ΔASA), number of intra and inter hydrogen bonds, salt bridges, Van-der-Waals (vdW) energy of different protease/BPTI complexes and interface sequence identity of proteases relative to BT.

In attempt to explain drastic differences in binding affinities of the three homologous PPIs, we explored ΔΔG_bind_ values between the three proteases and all single and double binding interface mutants of BPTI. To measure ΔΔG_bind_ values for tens of thousands of mutants in these three PPIs, we employed a recently developed by our group strategy that relies on yeast surface display (YSD) technology, NGS analysis and subsequent normalization of NGS data using a small dataset of ΔΔG_bind_ values measured of purified proteins (45). Applying such a protocol to the BPTI/BT interaction, yielded a very high (R > 0.9) correlation between NGS-based and actual experimental values of ΔΔG_bind_ and allowed us to predict the effect of all single mutations in the BPTI/BT interaction on the PPI binding affinity (45). In the present study, we extended the above approach to quantify ΔΔG_bind_ values for single and double mutants of BPTI interacting with CT and MT and for double mutants of BPTI interacting with BT.

Our data demonstrate that the three homologous BPTI/protease complexes possess drastically different binding landscapes and lie at different points in respect to the binding landscape maximum. Additionally, these differences in landscape contour and placement underlie correspondingly different energetic consequences of mutation, including asymmetrical directionality and different tendencies toward positive or negative epistasis.

## Results

To map binding landscapes of the three homologous BPTI/protease complexes, we first incorporated the wild-type BPTI (BPTI_WT_) gene into the pCTCON vector, compatible with YSD technology. In such a construct, BPTI_WT_ is expressed on the surface of a yeast cell with a C-terminal myc-tag for monitoring protein expression (Figure 2A). Binding of a protease to BPTI_WT_ was accessed by monitoring fluorescence of the FITC fluorophore conjugated to the protease via neutravidin. The assessment of binding of BPTI_WT_ to the three proteases by Fluorescently Activated Cell Sorting (FACS) showed a diagonal narrow distribution, demonstrating that BPTI_WT_ is well expressed on the surface of yeast cells, is properly folded, and binds to each of the proteases (Figure S1).

**Figure 2:**
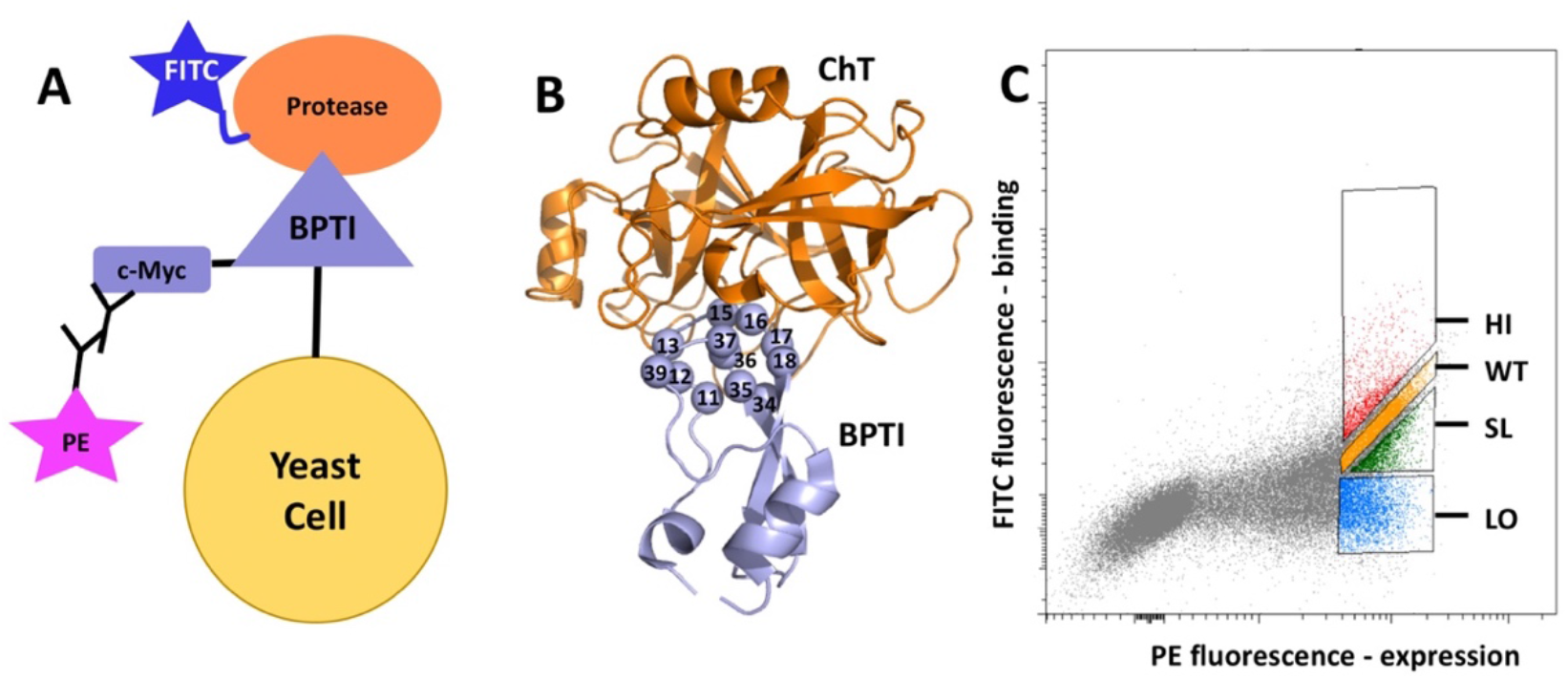
A) Yeast surface display construct with BPTI displayed on the surface of yeast cells. C-Myc tag is used to monitor BPTI mutant expression using PE-labeled antibody. Proteases are labeled by FITC that is used to monitor binding between the two proteins B) Construction of the BPTI mutant library. Structure of the ChT/BPTI complex with ChT shown in orange and BPTI in violet. BPTI binding interface positions randomized to twenty amino acids are shown as spheres (C) FACS data showing sorting of BPTI mutants binding to protease into four different populations. The uppermost HI gate contains BPTI mutants with affinity higher than that of BPTI_WT_. The second uppermost gate, WT, contains BPTI mutants with an affinity similar to BPTI_WT_. The third gate, SL, contains BPTI mutants with an affinity slightly lower than that of BPTI_WT_ and the lowest gate, LO, contains BPTI mutants with an affinity much lower than that of BPTI_WT_, The data is shown for the BPTI/ChT interaction, while similar data was obtained for the BPTI/MT and BPTI/BT interactions.

We next generated a library of BPTI mutants that contained all single and double BPTI mutants at positions that comprise the direct binding interface with proteases in the BPTI/protease structures. We randomized twelve BPTI positions to twenty amino acids while leaving two cysteines that participate in a disulfide bond intact to preserve BPTI folding (Figure 2B). In addition, all possible combinations of double mutations encompassing these twelve positions were encoded in the library. The BPTI library, referred to as the naïve library, contained 228 single mutants and all possible pairs of such mutations, resulting in the total theoretical diversity of 26,400 BPTI sequences. The naïve library was transformed into yeast and sequenced by NGS. Sequencing results showed that all possible single mutations and 89% of all double mutations were covered in the naïve library (60% when a cutoff of 5 sequencing reads was applied).

We next expressed the BPTI library on the yeast surface and measured expression and binding of the BPTI library to the three proteases using FACS (Figure 2C). Concentration of each protease was optimized to exhibit a considerable spread of the FACS binding signals from different BPTI mutants (Supplementary Figure S2). For each protease, we performed a sorting experiment and collected yeast cells with BPTI mutants belonging to four different affinity groups: higher than WT affinity (HI), WT-like affinity (WT), slightly lower than WT affinity (SL), and strongly lower than WT affinity (LO) (Figure 2C and Supplementary Figure S3-S5). The cells from each affinity gate were grown and sequenced with NGS, resulting in 300-900K reads per each population. For each BPTI mutant and each protease, we next calculated the enrichment value, which represents the ratio between the mutant’s frequency in a particular affinity gate to its frequency in the naïve library. We thereby obtained heatmaps of the enrichment values for all positions as shown on Figure 3 for the CT/BPTI interaction.

**Figure 3:**
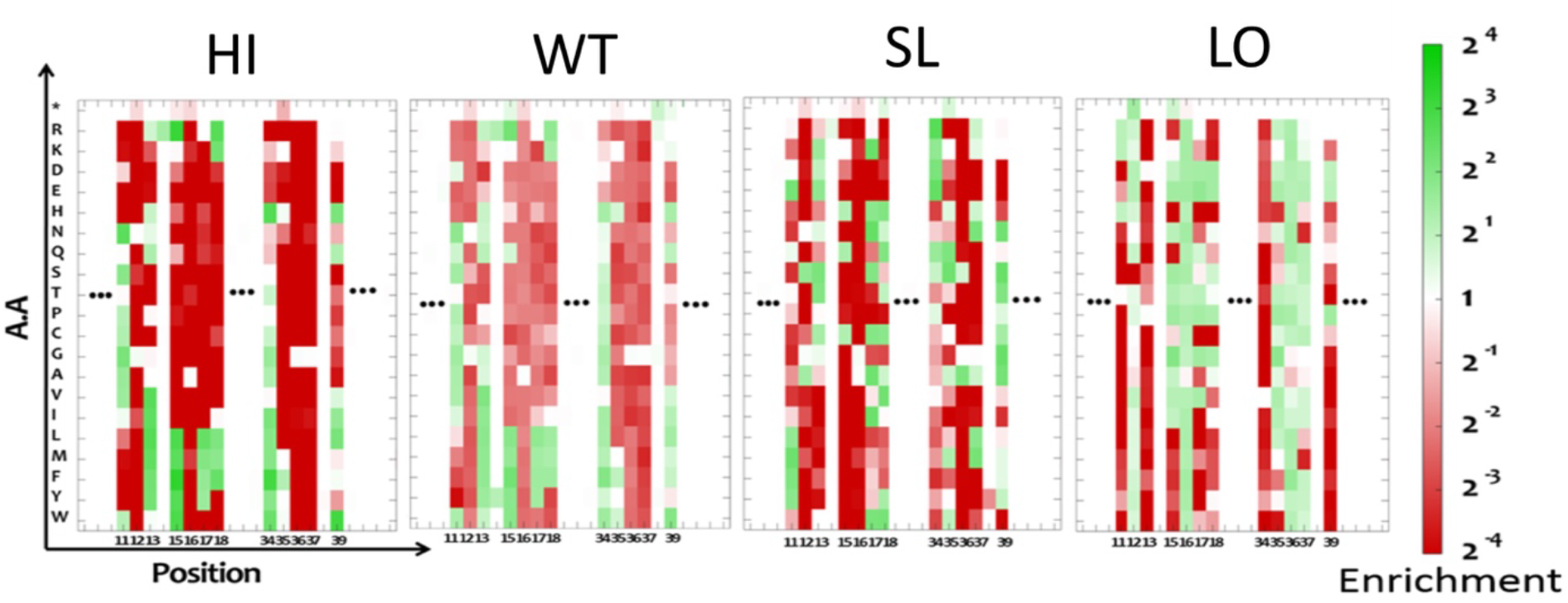
Heatmap showing the enrichment values for each single mutation in the ChT/BPTI complex in the four sorted affinity gates. The enrichment ratio varies from high (green) to low (red) as shown on the right axis. Similar maps were obtained for the other two PPIs.

While the enrichment maps give us qualitative measures of affinity changes due to various mutations, our goal was to construct and compare quantitative binding landscapes of the BPTI/protease interactions. We thus utilized the methodology developed in our recent paper that allows us to normalize the NGS-based enrichments using a small dataset of experimental ΔΔG_bind_ values measured by biophysical techniques on purified proteins (45). We first compiled such normalization datasets for the three complexes, collecting 34 and 33 ΔΔG_bind_ data points from literature for the ChT/BPTI and BT/BPTI interactions, respectively (38, 45–48). For the MT/BPTI interaction, where only a few ΔΔG_bind_ data points have been reported (44, 49), we produced the normalization dataset by expressing and purifying 12 BPTI mutants and measuring their binding affinities to MT (Figure S6). The above datasets were used to obtain a normalization formula for each protease that converts the four enrichment values from the NGS data into the predicted ΔΔG_bind_ values. For all three enzymes, high correlation was found between the ΔΔG_bind_ values predicted from NGS and those experimentally determined using purified proteins (R= ~0.9; Figure S7).

We next used the acquired normalization formulas to predict ΔΔG_bind_ values for all single and double BPTI mutants that have been detected by NGS for the three PPIs. While nearly all BPTI single mutants have been sequenced in all four affinity gates for the three proteases, the double mutants were covered less extensively in the NGS results with only 576, 3393, and 636 double mutants appearing in all four affinity gates for ChT, BT and MT, respectively. To increase the coverage of ΔΔG_bind_ predictions for the double mutants and to complete the predictions for single mutants, we examined whether normalization formulas could be obtained from subsets of three, two, and one affinity gates. While all subsets of gates were examined, only those subsets that produced high correlation with experimental data on pure proteins were selected for the final predictions. For each ΔΔG_bind_ prediction, we estimated the uncertainty in ΔΔG_bind_ predictions using the bootstrapping of the NGS data as described in detail in our previous work (45). Overall, we were able to make reliable predictions for 13,113 double mutants for the BT/BPTI interaction (50% of all binding interface double mutations), 12,537 for ChT/BPTI interaction (47%), and 4317 for MT/BPTI interaction (16 %). We thus constructed full single-mutant binding landscapes and partial-double mutant binding landscapes for BPTI interacting with the three homologous proteases with highly divergent K_D_s.

### Analysis of the single-mutant binding landscapes

To compare how single mutations affect free energy of binding in the 3 PPIs, we summarized our results in a histogram that includes ΔΔG_bind_ values from all 228 single mutations for each PPI (Figure 4A-C). While all the three histograms show predominance of destabilizing mutations (ΔΔG_bind_> 0), the magnitude of destabilization due to single mutations differs substantially among the three PPIs. For the high-affinity BT/BPTI complex, very high ~12 kcal/mol destabilizations were observed due to some single mutations, medium destabilizations (up to 6 kcal/mol) were observed in the ChT/BPTI complex and small destabilizations (up to ~3 kcal/mol) were observed for the low-affinity MT/BPTI complex (Figure 4 and Figure S8). On average, a single mutation destabilized BT/BPTI interaction by 4.5 kcal/mol, ChT/BPTI interaction by 1.6 kcal/mol and MT/BPTI interaction by 0.82 kcal/mol. On the contrary, affinity-enhancing mutations appeared more frequently in the low-affinity MT/BPTI complex (50 mutations or 22%), less frequently in the medium-affinity ChT/BPTI complex (37 mutations or 16%), and only once (<1%) in the high-affinity BT/BPTI complex. Per-position analysis of ΔΔG_bind_ values revealed that all but one position on BPTI are absolute hot spots in the BT/BPTI interaction, exhibiting only positive ΔΔG_bind_ values. In contrast, only four absolute hot-spots are present in the ChT/BPTI interaction (positions 12, 16, 36, 37) and only two in the MT/BPTI interaction (position 16 and 36). The spatial distribution of cold-spot and hot-spot positions showed different patterns among the three complexes (Figure 5). Interestingly, position 15, which is central to the binding interface, is a cold spot in the ChT/BPTI interaction with all hydrophobic amino acids leading to improved affinity. The same position contains highly destabilizing mutations in both the BT/BPTI and the MT/BPTI complexes, except for one mutation K15R that leads to affinity improvement in both complexes. These differences in the position 15 preferences are in complete agreement with previous studies on purified proteins for the BT/BPTI and ChT/BPTI complexes (47). Additionally, amino acid preferences at position 15 of BPTI discovered here reflect the preferences for substrates that these enzymes cleave (Lys and Arg for trypsins and hydrophobic amino acids for chymotrypsins), indicating that these enzymes have evolved to possess optimal binding pockets for these amino acids.

**Figure 4:**
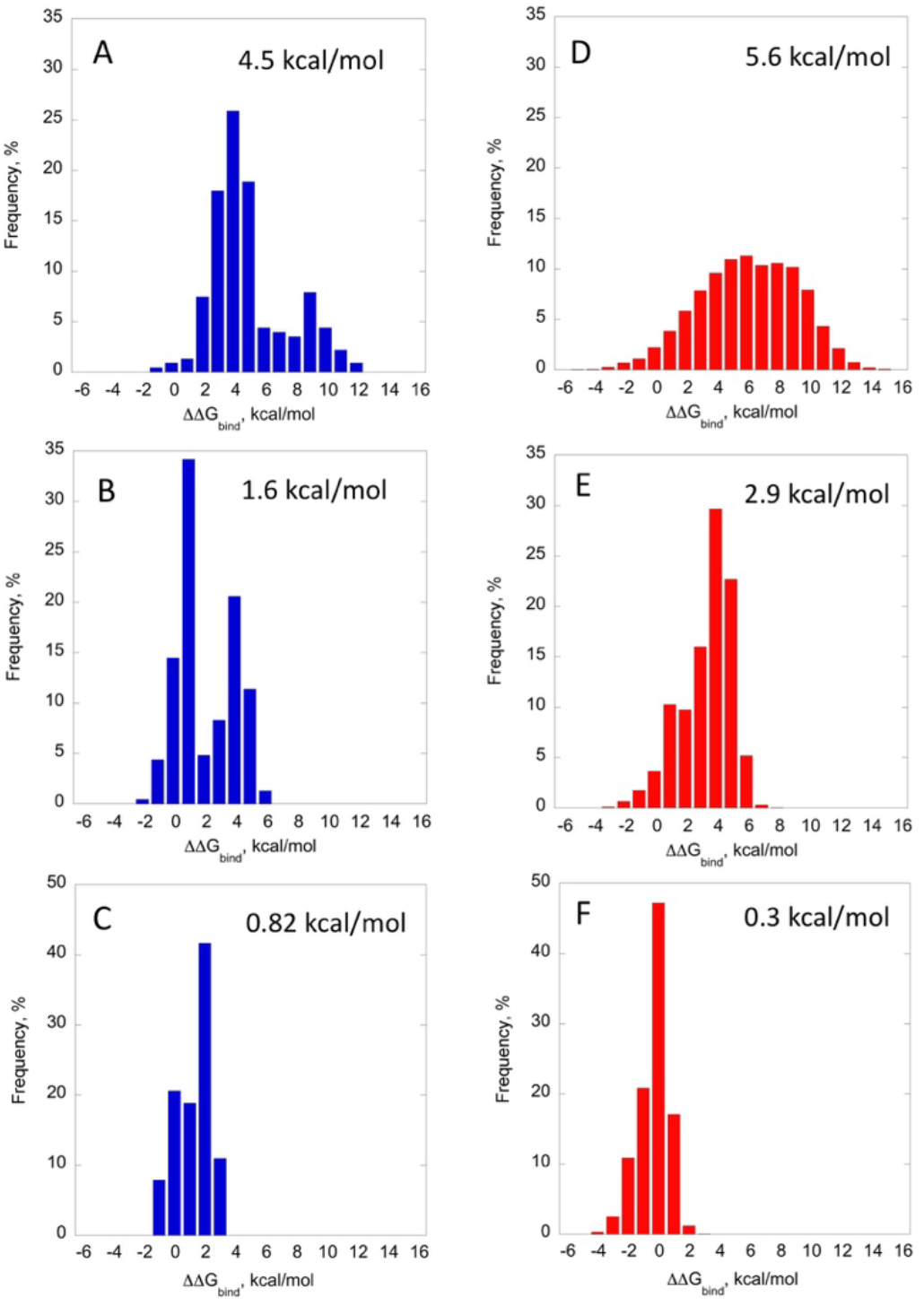
Histograms of ΔΔG_bind_ for single and double BPTI mutants. Single BPTI mutants interacting with (A) BT; (B) ChT; (C) MT. Double BPTI mutants interacting with (D) BT; (E) ChT; (F) MT. Mean value for ΔΔG_bind_ for each histogram is displayed on top of each graph. While all 228 single mutants are incorporated into the single-mutant histograms for all proteases; only ~5O% of double mutants are summarized for BT and ChT and only 16% for MT. The data is available in the Source_data_file.

**Figure 5:**
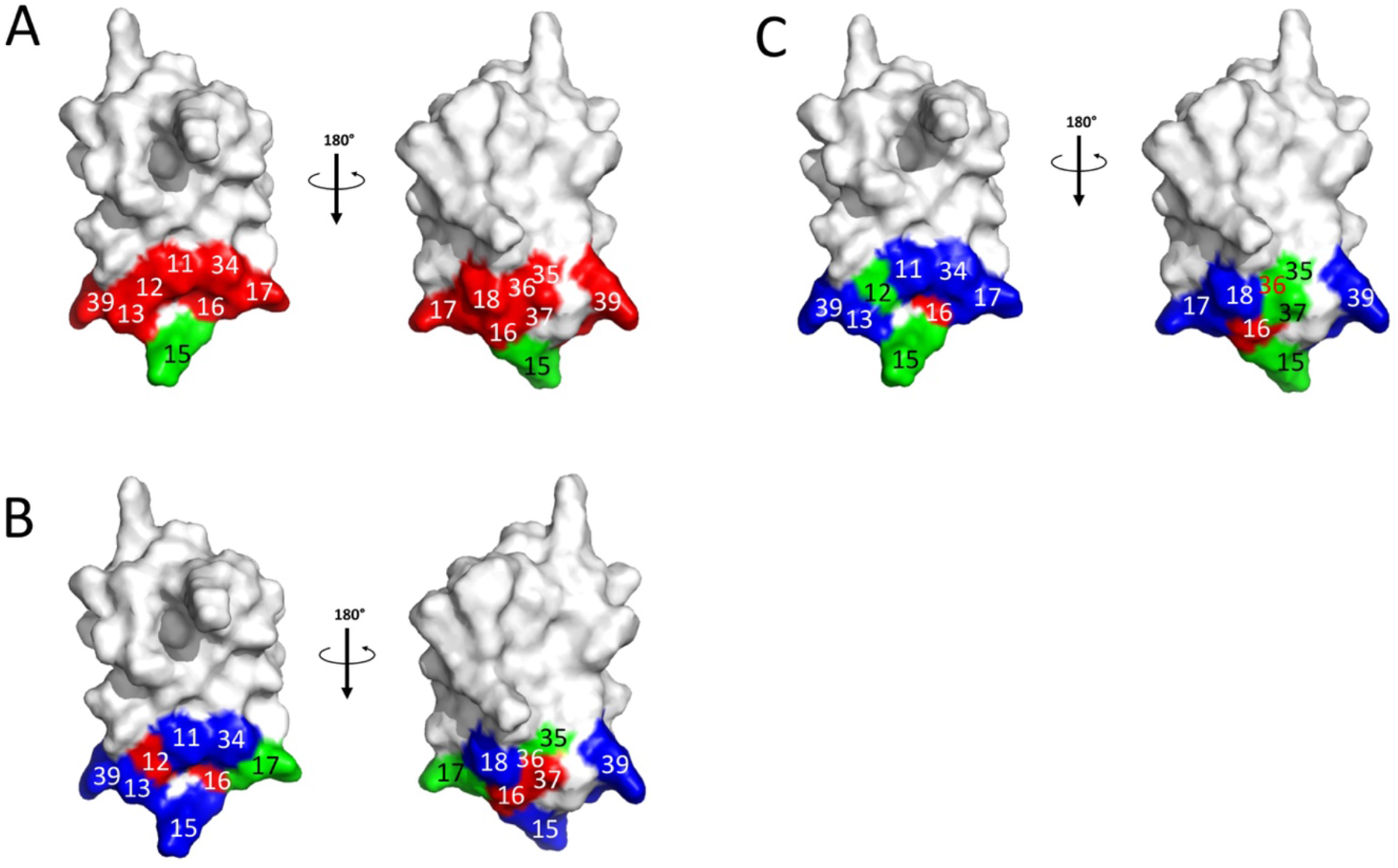
Structure of BPTI with binding interface positions colored according to the number of detected affinity-enhancing mutations at this position when interacting with (A) BT; (B) ChT; and (C) MT. Red - no affinity-enhancing mutation was detected; green - 1-2 affinity-enhancing mutations were detected; blue - 3 or more affinity-enhancing mutations were detected.

### Analysis of the double-mutant binding landscapes

We next compared the double-mutant binding landscapes for the three PPIs. We first plotted the histograms of ΔΔG_bind_ for the double mutations, for which ΔΔG_bind_ predictions were available (Figure 4D-E). Our results show that on average a double mutation destabilizes the high-affinity BT/BPTI complex by 5.9 kcal/mol, the medium affinity ChT/BPTI complex by 2.9 kcal/mol and the low-affinity MT/BPTI complex by 0.3 kcal/mol, showing the same tendency of increased destabilization due to double mutation with increasing affinity of the PPI as was observed for single mutants. When comparing an average effect from a double and a single mutation, BT/BPTI and ChT/BPTI exhibited higher ΔΔG_bind_ value for a double mutation, consistent with a majority of single mutations being destabilizing in these two PPIs. For the low-affinity MT/BPTI complex, the double mutant average was slightly lower compared to that of the single mutant average. This could be due to the fact that a large fraction of single mutants for this PPI leads to affinity improvement or due to the relatively small coverage of double mutants for this PPI in our study (only 16% of double mutations had ΔΔG_bind_ predicted).

Using the extensive ΔΔG_bind_ data for double mutations, we further explored how a single mutational step from a WT sequence alters the distribution of ΔΔG_bind_ values for the second mutation. For this analysis, we selected three representative single BPTI mutants in the highest-affinity BPTI/BT complex: BPTI_K15R that shows slight improvement in affinity compared to BPTI_WT_ (ΔΔG_bind_= −1.4 kcal/mol), BPTI_A16S whose affinity to BT is considerably weaker in comparison to BPTI_WT_ (ΔΔG_bind_= +4.5 kcal/mol) and BPTI_K15A that shows dramatically reduced affinity in comparison to BPTI_WT_ (ΔΔG_bind_= +11.1 kcal/mol). We next compared the ΔΔG_bind_ distributions for single mutations taken on the background of each of the three specified first mutations. While only partial single-mutant landscapes could be constructed for these three BPTI mutants interacting with BT (as we have the data for ~50% of the double mutants), for the detected mutants we observe significant differences in the binding landscapes of the three BPTI mutants with BT ion (Figure S9). K15R that improves the fitness of the BT/BPTI interaction produces a histogram with mostly destabilizing mutations going as far as +12 kcal/mol, yet some affinity-enhancing mutations are also observed. The medium-destabilizing mutation A16S results in a landscape that contains both stabilizing and destabilizing steps with magnitudes ranging from −6 to +8 kcal/mol. The highly destabilizing mutation K15A exhibits a landscape that mostly contains stabilizing mutations with the highest stabilization of −6 kcal/mol. Note that for the K15A mutant we do not observe a mutational step that would reach the affinity of the WT BT/BPTI complex. This is likely due to the fact that position 15 is the most important energetically for the BPTI/protease interaction, thus destroying the favorable interaction at this position could not be fully compensated by any other mutation on BPTI. Our results hence indicate that with every mutational step taken from the WT BPTI sequence the binding landscape would be changed depending on the first mutation; this change is a result of non-additivity of some of single mutations in BPTI.

Next, using the extensive quantitative data on ΔΔG_bind_ for single and double mutants, we investigated the extent of coupling between various point mutations in BPTI when it interacts with the three proteases. We have classified mutations into three classes: additive, exhibiting positive and negative epistasis according to the magnitude of the coupling energy *ΔGi* upon two mutations X and Y:

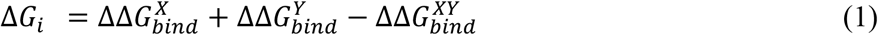

Here, 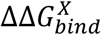 and 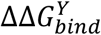 represent the change of the binding free energy of the single mutants X and Y, respectively, 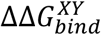 represents the change of binding free energy of the double mutant containing mutations X and Y. Two mutations were defined as additive if △*G_i_* is zero within the uncertainty of our ΔΔG_bind_ predictions (see Methods for details). Positive epistasis was defined when Δ*G_i_* >0, i.e the double mutation exhibits higher fitness compared to what is expected from additivity of two single mutations. Negative epistasis was defined when Δ*G_i_* <0, i.e the double mutation exhibits lower fitness compared to what is expected from additivity of the two single mutations.

Coupling energy analysis shows that in the BT/BPTI interaction, 59% of the detected mutations are additive, 40% of mutations show positive epistasis, and only ~1% of mutations show negative epistasis. In the ChT/BPTI interaction, 74% of mutations are additive, 18% of mutations show positive epistasis and 8% of mutations show negative epistasis. Finally, among the detected mutations in the MT/BPTI interaction we observe 70% of mutations are additive, 26% exhibit positive epistasis and 4% - negative epistasis. Note that the smaller fraction of mutations with negative vs. positive epistasis observed for all complexes could be in part due to the fact that we do not see highly destabilizing double mutations in our experiment and such mutations are likely to be among those exhibiting negative epistasis. We further constructed the per-position correlation matrices displaying coupling energy between all detected mutations in the three PPIs (Figure 6). Figure 6 shows that the sign of the epistasis depends not only on a pair of positions but also on the mutation type. Yet, certain preference for either negative or positive epistasis frequently dominates coupling at certain position pairs as some squares are predominantly red or blue on Figure 6. To analyze which positions exhibit higher degree of coupling, we averaged ΔGi values over all detected mutations at each pair of positions (Figure 7). Figure 7 shows that different proteases exhibit different patterns of coupling between pairs of mutations. Interestingly, positions where highest destabilization is observed for single mutations show high degree of positive epistasis with all other positions (such as for example positions 15 and 16 in the BPTI/BT complex, position 12 in the ChT/BPTI complex and positions 35, 36 in the MT/BPTI complex). On the other hand, cold spot positions tend to exhibit negative epistasis with many other positions in the protein (see for example, positions 13, 18, 34, 39 in the ChT/BPTI complex and 34 in the MT/BPTI complex). The average coupling energies are larger for the highest affinity BT/BPTI complex, medium for the CT/BPTI complex and the lowest for the MT/BPTI complex (Figure S10). We further tested whether the degree of coupling between the two mutations depends on the distance between the mutated positions (Figure S11). Our results show that mutations at two closely-located positions could exhibit various degrees of coupling from high to low. As the distance is increased between positions, the average coupling between the two mutations at these positions decreases (Figure S11). A similar trend has been observed for all three PPIs and is in agreement with previous studies in various biological systems (50).

**Figure 6:**
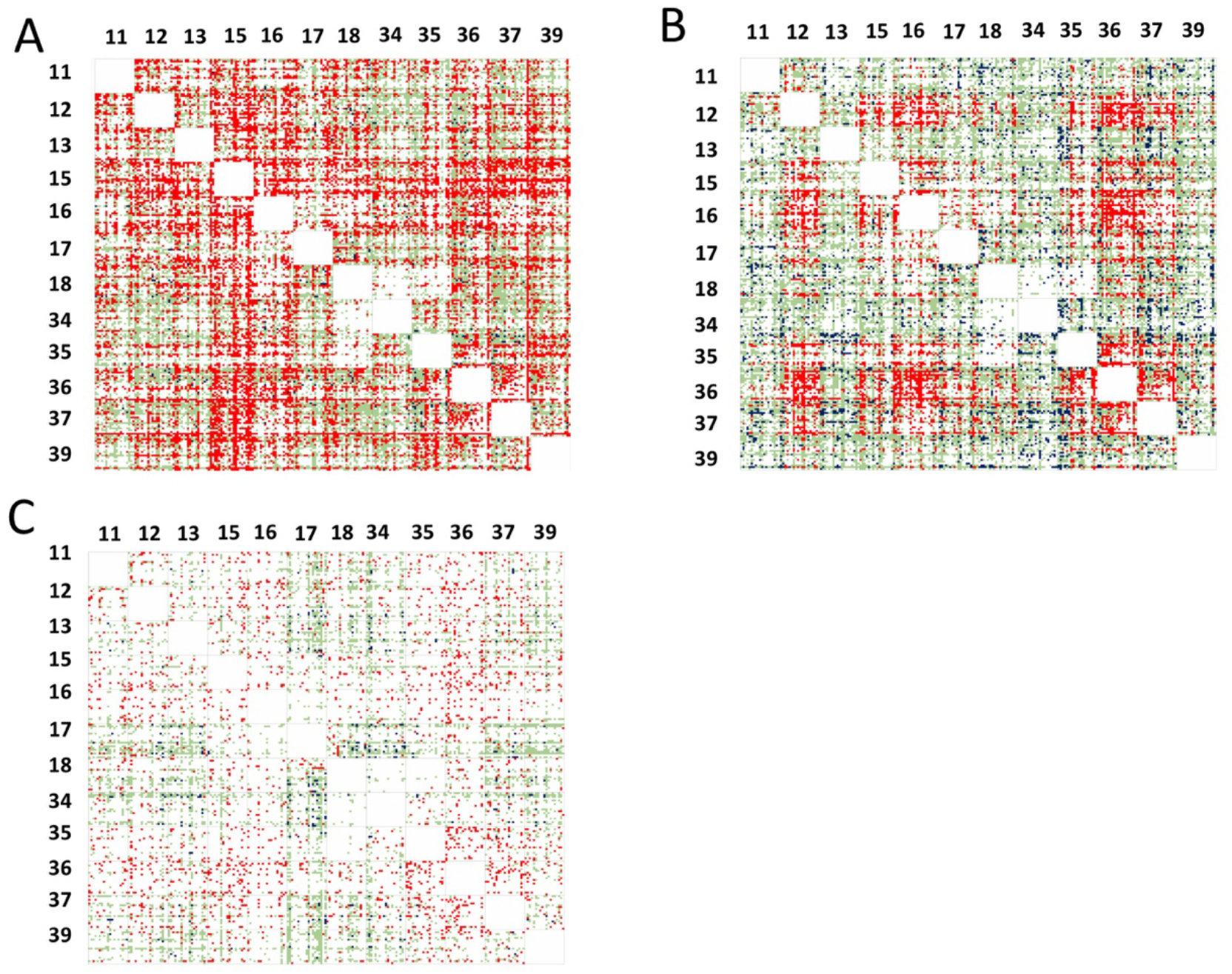
The matrix of ΔGi for double mutations in the three PPIs: (A) BT/BPTI; (B) ChT/BPTI and (C) MT/BPTI. On the left and on the top are BPTI binding interface positions randomized to 20 amino acids. Each square is a 20×20 matrix containing ΔGi values for coupling between a particular mutation at one position to another particular mutation at another position in the same order. Color coding shows the degree of cooperativity: Green: additivity; red - Positive Epistasis; Blue-Negative Epistasis; white - no data is available.

**Figure 7:**
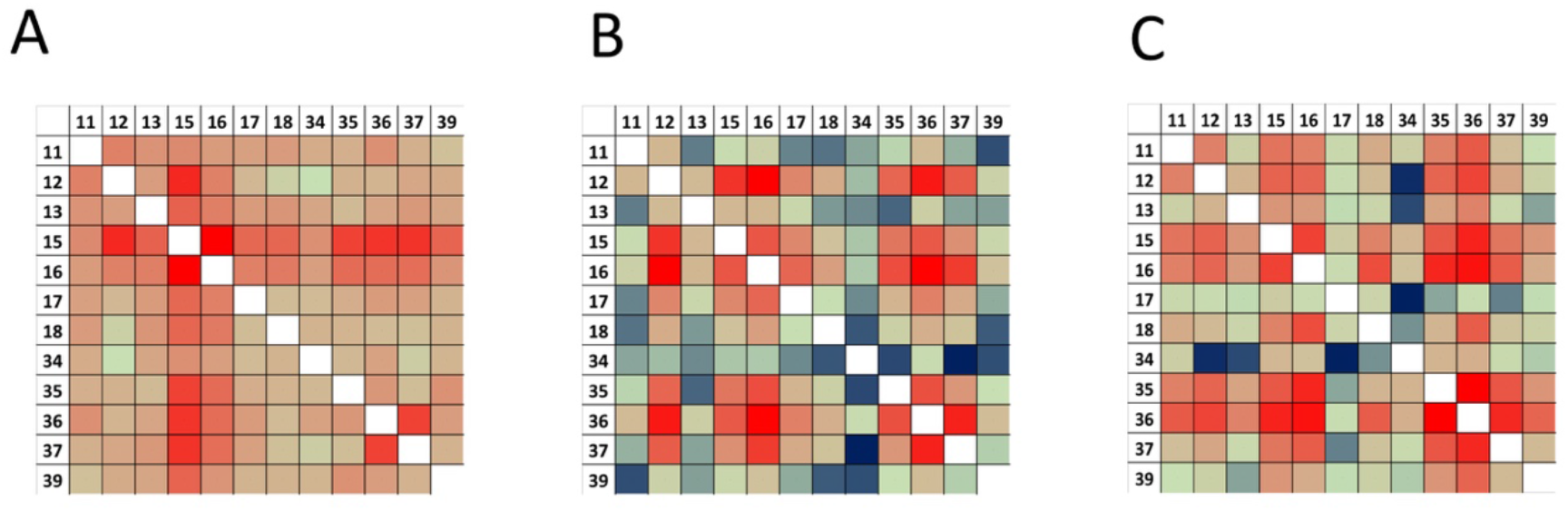
Coupling energies *ΔGi* averaged over all detected mutations at a pair of BPTI positions: (A) BT/BPTI interaction; (B) ChT/BPTI interaction; (C) MT/BPTI interaction. On the top and on the left BPTI positions are shown. The values are color coded from high positive epistasis (dark red) to additive (green) and to negative epistasis (dark blue).

## Discussion

In this study we measured quantitative effects of tens of thousands of single and double mutational steps in three homologous enzyme-inhibitor complexes. While the complexes are similar in their sequences and structures, they differ greatly in binding affinities that range from ultra-high to low. We find that the binding landscape of each PPI depends strongly on the interaction K_D_. In particular, the ultra-high affinity BT/BPTI complex is highly evolutionary optimized. Accordingly, the sequence of WT BPTI lies nearly at the maximum of the binding landscape, with only one mutation leading to significant affinity improvement. The landscape also exhibits a steep gradient, with a majority of single mutations leading to large steps down the hill with a maximum drop of ~12 kcal/mol and an average drop of 4.5 kcal/mol (45) (Figure 8). Such high destabilizations from single mutations are extremely rare. For example, the SKEMPI database (51) that reports 5079 single mutant binding affinity changes in various PPIs contains only 16 single mutations (0.3%) with ΔΔG_bind_ values greater than 8 kcal/mol, and all of them belong to high-affinity complexes (with a K_D_ of 10^-12^ M or better). The medium-affinity ChT/BPTI complex shows lower degree of evolutionary optimality, with a larger fraction of mutations leading to affinity improvement (16%) and a maximum improvement of 2.6 kcal/mol. Yet even in this complex, single mutational steps could lead to high complex destabilization of up to 6 kcal/mol. Thus, the landscape of the ChT/BPTI complex exhibits a medium gradient and the WT sequence lies about 2/3 up the landscape hill (Figure 8). One might expect that the low-affinity MT/BPTI complex would exhibit ΔΔG_bind_ distribution that is the opposite of that observed for the BPTI/BT complex, with high number of mutations that lead to very large improvement in binding affinity. Yet, this is not what we observe in the present study. The MT/BPTI complex indeed exhibits the highest fraction of mutations leading to affinity improvement among the three complexes (22%) but the largest improvement due to a single mutation does not exceed 1.9 kcal/mol, smaller than what is observed for the ChT/BPTI complex. Yet, the reduction in binding affinity due to single mutations is also the smallest for the MT/BPTI complex not exceeding 3 kcal/mol. Thus, we conclude that the difference between the MT/BPTI and BT/BPTI complexes is not only the location of their sequences relative to the maximum of the binding landscape, but the landscapes themselves show different gradients, high for the high-affinity complex and low for the low-affinity complex (Figure 8).

**Figure 8:**
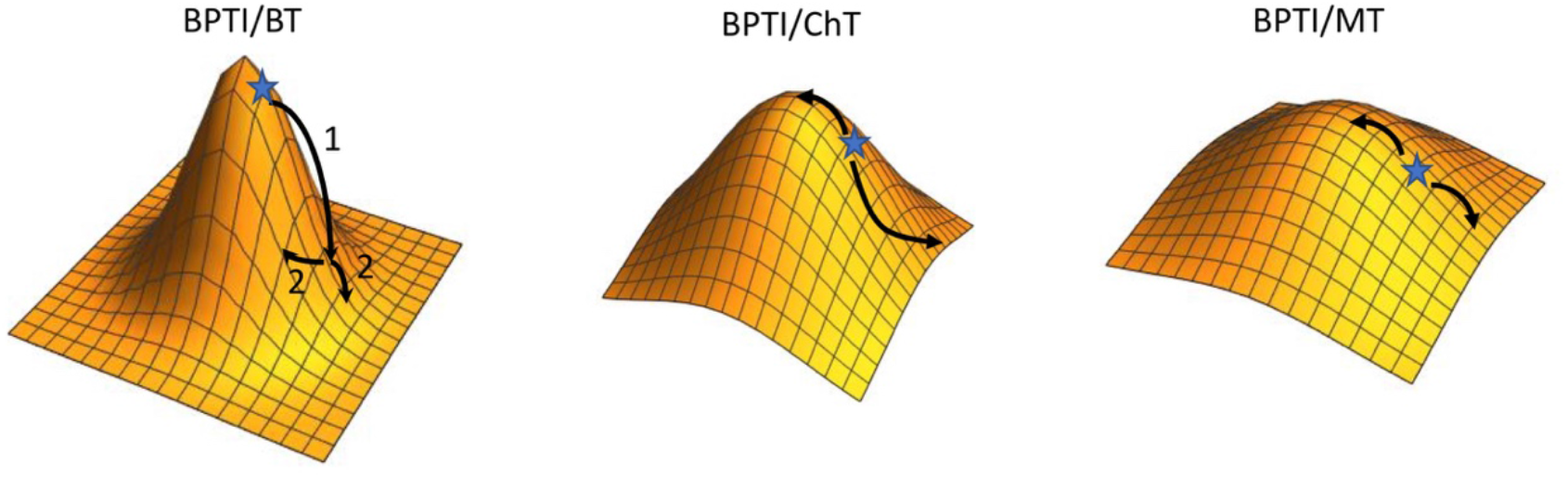
Schematic illustration of the single-mutant binding landscapes for the three studied PPIs. The maximum on the surface corresponds to the highest possible binding affinity. A star indicates the position of the WT BPTI sequence in respect to the maximum. Arrows illustrate how single mutations lead to affinity changes in the three complexes.

The binding landscape characteristics of the three studied PPIs have been dictated by their evolutionary history. BPTI is a cognate inhibitor of BT, thus both proteins have coevolved in one organism to optimize their affinity for each other. ChT and MT bind to BPTI only due to homology to BT. ChT has other cognate inhibitors with which it interacts with much higher affinity, such as turkey ovomucoid third domain (OMTKY3) (K_D_ of 1.9×10^-11^) (52). MT, by contrast, appears to have evolved for widespread natural resistance to the tight-binding mechanism of the proteinaceous canonical trypsin inhibitors such as BPTI (53),(54). Furthermore, by binding to these inhibitors orders of magnitude more weakly than other trypsins, mesotrypsin has evolved the capability to cleave many proteinaceous trypsin inhibitors as substrates (55–59). This functional role of the MT/BPTI interaction agrees with our finding that no single or double mutation on the BPTI side could convert this complex into a high-affinity PPI.

### Analysis of cold spots

In this study we identified a number of cold spot positions in ChT/BPTI and MT/BPTI complexes at which multiple mutations lead to affinity improvement. In our previous study we identified two scenarios of how cold spots occur either through removal of unfavorable interaction with the partner protein or through introduction of a new favorable interaction, where no interaction exists (43). Cold spots discovered here could be easily fit into these two scenarios. We observe the first scenario occurring in the MT/BPTI complex at position 17, where an Arg on BPTI is found in close proximity to an Arg 193 on MT. Substituting Arg 17 with a small and/or hydrophobic amino acid results in affinity improvement. On the contrary, Arg 17 of BPTI is found in a largely hydrophobic environment in the BPTI/ChT complex; its replacement with a hydrophobic Met and Leu results in slightly negative ΔΔG_bind_ values.

We observe the second scenario for cold spot formation at position 34 in the ChT/BPTI and MT/BPTI complexes. V34 at this position does not form any interactions with these proteases. Its replacement with larger hydrophobic amino acids that bury additional surface area increases affinity to ChT. Its replacement with polar or negatively charged residues improves affinity to MT by likely forming new hydrogen bonds to Tyr 151 and/or Gln 192 on MT.

### Epistasis in protease/BPTI complexes

Using the data for tens of thousands of double mutants we were able to analyze how two mutations are coupled in the three protease/BPTI complexes. Our data shows that in the high-affinity BT/BPTI complex a large proportion of double mutations results in positive epistasis and only a minority of mutations produces negative epistasis. More equal distribution of mutations with positive and negative epistasis is observed for the ChT/BPTI complex. Note, that we only observe ~50% of all double mutations for these two PPIs; the remaining 50% that are invisible in our experiment are likely to be highly destabilizing, which would put them in the category of either additive or showing negative epistasis. The abundance of positive epistasis in the BT/BPTI interaction could be explained from the perspective of binding landscape theory (Figure 8). Due to the steepness of the gradient in the area of the wild-type BPTI sequence, the first mutation in this PPI leads to a large step down the hill into the area of low-gradient. Second mutation from this point could lead up or down, but the change would be relatively small, resulting in positive epistasis, i.e., better ΔΔG_bind_ compared to what would be predicted from additivity of the two highly destabilizing mutations. Positive epistasis particularly predominates at positions where the largest affinity drops are recorded (such as at positions 15 and 16 for the BT/BPTI complex or position 12 for the ChT/BPTI complex), where the gradient is steep.

Positive epistasis could be also explained from the structural perspective. Highly optimized PPIs usually retain their original binding conformation upon introduction of a single mutation due to the abundance of favorable interactions generated at non-mutated positions. Yet, if any of the hot-spot residues is mutated, substantially weakening the interaction, then the impact of a second deleterious mutation may be mitigated by an increase in flexibility at the interface, enabling the protein to adopt alternative conformations that introduce new favorable intermolecular contacts and enhance affinity. If the same mutation would occur on the background of the wild-type residue in the hot-spot position, the new conformation would not be accessible and the affinity enhancement would not be achieved. Thus, such a double mutant would possess better ΔΔG_bind_ compared to the sum of single mutants, exhibiting positive epistasis.

On the contrary, negative epistasis is more frequent for medium and low-affinity PPIs and appears mostly when one mutation is performed at a cold-spot position. If the first mutation improves binding affinity and thus makes a step up the binding landscape towards the maximum, the second mutation would be made from the point of steeper gradient and is likely to make a large step down, thus resulting in negative epistasis. Structurally that means that if at one cold spot position a new favorable interaction was created, this interaction might lock the PPI into a new slightly different conformation. Another conformation might be acquired upon introduction of a different affinity-enhancing mutation. But the two favorable conformations could not be achieved simultaneously, resulting in worse ΔΔG_bind_ for a double mutation compared to the sum of two single mutations (negative epistasis).

In summary, in this study we report ΔΔG_bind_ values for tens of thousands of single and double mutations in three protease/BPTI complexes with similar structures but highly variable binding affinities, thus generating an unprecedented amount of mutational data that could be used as benchmark for testing new computational methodology and for design of new high-affinity protease inhibitors. Using the obtained data, we demonstrate striking differences between the binding landscapes of the three PPIs that could be explained by the level of the PPI evolutionary optimality. Furthermore, we study how two single mutations in these PPIs couple to each other and demonstrate that the coupling energy depends not only on positions of mutations but also on the identities of the mutated amino acid. Furthermore, we observe that mutations at hot-spot positions generally exhibit positive epistasis with other mutations while mutations at cold-spot positions generally exhibit negative epistasis and explain this phenomenon from the perspective of binding landscape theory. Our powerful experimental methodology could be used to access the binding landscapes in many additional PPIs with different structures, functions, and affinities and to probe whether the reported evolutionary trends hold in other biological systems.

## Supporting information

Dataset S1 contains all ddGbind values

## Acknowledgements

We thank U. Hadad (BGU) for help with FACS experiments. This work was supported by the Israel Science Foundation (ISF) grant 1873/15 (J. M. S.) and by the European Research Council (ERC) grant 336041 and the ISF grant 1615/19 (N. P.). N. P. and E. S. R. acknowledge support from the US-Israel Binational Science Foundation (BSF). J. S. M. acknowledges support from the US-Israel Binational Science Foundation (BSF) and the Israel Cancer Research Foundation (ICRF). E. S. R. acknowledges support from U.S. National Institutes of Health grants R01 CA154387 and R01 GM132100.

## Author contributions

J. M. S. and N. P. designed the research. M. H., J. S. and I. C. performed the research. M. H. and J. M. S. performed data analysis. Y. P. provided help with combinatorial library construction and transformation. E. S. R. provided protease samples. J. M. S. wrote the manuscript with assistance from N. P. and E. S. R.

## Methods

### BPTI library construction

Twelve positions on BPTI that lie in the binding interface with BT (PDB ID 3OTJ) were subject to randomization: T11, G12, P13, K15, A16, R17, I18, V34, Y35, G36, G37 and R39. A BPTI library was constructed that randomized two positions at a time with an NNS codon (where N = A/C/G/T DNA base, S = C/G DNA base); encoding all amino acids at the randomized positions, including the WT amino acid. The library was divided into 66 sublibraries that each incorporates all possible pairs of the twelve randomized positions. TPCR protocol(60) was used to create each library using two primers that either combined two mutations in one primer or divided them into two primers depending on their proximity to each other (Supplementary Note 1 in Supplementary information). These primers were used in a PCR together with BPTI_WT_ plasmid to incorporate these mutations at the specific positions in BPTI and to amplify the whole plasmid. Agarose gel analysis was used to confirm the success of each TPCR reaction. The TPCR-amplified plasmid DNA was treated with DpnI (New England Biolabs, Ipswich, MA) to remove any parental plasmid used as a template to construct the library, cleaned up with magnetic beads (AMPure XP, Beckman coulter, Brea, CA), transferred into *E. coli* and selected colonies were sequenced to confirm the successful generation and transformation of the BPTI library. The vectors containing the BPTI library were extracted using QIAprep Spin miniprep (Qiagen, Hilden, Germany) and all the sub-libraries were pooled together and balanced by their DNA amount to use the same amount of DNA from each sub-library (~3.6μg). Then, the pooled library was transferred into *S. cerevisiae* using 20 transformations resulting in 60,000 – 70,000 colonies for the complete library as estimated by plating 1/20 amount of the library sample and counting the colonies after transformation on a SDCAA plate.

### NGS analysis

The paired-end reads from the NGS experiments were merged(61) and their quality scores were calculated in the FastQC tool (https://www.bioinformatics.babraham.ac.uk/projects/fastqc/). In the Matlab script, the sequences were aligned, and sequences containing extra mutations at non-randomized positions were filtered out. The number of each remaining BPTI mutant i was counted in the sorted and the naïve populations and its frequency f in the libraries was calculated (**Eq. 2**). Using the frequency of the mutant in one of the sorted populations and the naïve population, the enrichment e^i^ of each BPTI mutant was calculated (**Eq. 3**).

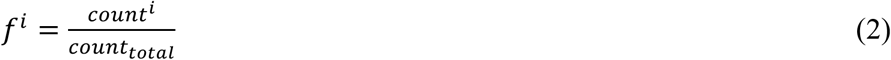

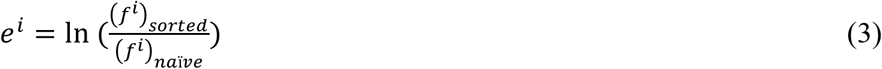

To estimate the uncertainty in BPTI mutant frequencies we applied a bootstrapping method to the NGS data for all sorted gates and the naïve library as described in(62). Briefly, the original NGS data was used to randomly draw sequences to obtain a resampling data set of the same size and to calculate the frequency of each BPTI mutant in each population. The resampling process was repeated 1000 times and the average frequency and the standard deviation was calculated from 1000 resampling data sets for each BPTI mutant in each sorting gate and in the naïve library. The error was propagated into equations 2 and 3 to calculate the error in enrichment values as described in(45).

### Predicting ΔΔG_bind_ values from NGS data

All available experimental data on ΔΔ*G*_bind_ for the BPTI/protease complexes was used to obtain the best normalization formulas for each complex for converting enrichment values from the four sorted populations into ΔΔ*G*_bind_ values. To this end, we used a linear regression model function in Mathematica (Wolfram Research) with 5 parameters (*Y*= *aX_1_*+*bX_2_*+*cX_3_*+*dX_4_*+f if all four enrichment values were available in our NGS data for this particular mutation. The parameters a, b, c, d, f were optimized using the experimental data set as values of Y and the set of *X_1_, X_2_, X_3_, X_4_* enrichment values. The obtained normalization formula (different for each protease) was used to calculate ΔΔ*G*_bind_ values for all the remaining single and double BPTI mutants that had four enrichment values recorded in the NGS experiment. To make ΔΔG_bind_ prediction for mutants where fewer than 4 enrichment values were available, we repeated the normalization procedure using different subsets of enrichment values (such as X_1_ and X_4_; X_1_, X_2_, X_3_; etc.). Accordingly, we varied the number of parameters in the normalization formula. We checked whether high correlation with the experimental data set of ΔΔG_bind_ values could be obtained using this particular subset of variables as predictors. If a correlation of R>0.80 was obtained between the predicted and the experimental ΔΔG_bind_ values, the set of gates was selected as good for making predictions. Additional cross check for validity of predictions from this subset of gates was performed by comparing ΔΔG_bind_ predictions for all single mutants based on all four gates and based on the selected subset of gates and confirming high correlation between the two predicted ΔΔG_bind_ values over all single mutations. For each of the mutant, we used the available enrichment values to make separate predictions from all possible “good” subset of gates. First all predictions were recorded for mutants where enrichment values were available for all 4 gates. For mutants where predictions were available for only gates X1, X2, and X4, predictions were made based on these 3 gates providing that this set of gates was defined as good. For mutants where predictions were available in only gates X1 and X4 predictions were made based on these two gates if this set of gates was defined as good for predictions. For each prediction from each subset of gates, the uncertainty of the prediction was calculated using the procedure outlined above. Finally, for each mutation, ΔΔG_bind_ prediction was selected from all the predictions according to the subset of gates where the highest correlation with experimental data was observed. The dataset of final ΔΔG_bind_ predictions for all single and double mutants for the three PPIs could be found in the Dataset S1 file.

### Production of BPTI variants

The BPTI_WT_ sequence was cloned into a pPIC9K vector (Invitrogen, Carlsbad, CA), TPCR was used for site-directed mutagenesis to create the sequences of all variants, transformed, expressed in *P. pastoris* (*GS115* strain; obtained from Invitrogen, Carlsbad, CA) and purified by nickel affinity chromatography, followed by size-exclusion chromatography, as described in a previous work^85^. The correct DNA sequence of each produced protein was confirmed by extracting the plasmidic DNA from *P. pastoris* after protein purification by nickel chromatography, amplifying the BPTI gene and sequencing it. Protein purity was validated by SDS-PAGE on a 20% polyacrylamide gel, and the mass was determined with a MALDI-TOF REFLEX-IV (Bruker) mass spectrometer (IKI, BGU; data not shown). Purification yields for all BPTI variants were 2-15 mg per liter of medium. The concentration of purified BPTI variants was determined by an activity assay.

For MT, values of the inhibition constant (*K_i_*) were determined using a general enzyme activity essay for PPIs characterized by medium to low affinity^62^. Here, 304 μL BPTI (4 different concentrations ranging from 5.2 to 52.6 μM) was mixed with 8 μL of the substrate Z-GPR-pNA (Sigma-Aldrich, St. Louis, MO) (5 different concentrations ranging from 0.4 to 10 mM). The mix was incubated for three minutes. Then, the reaction was initialized by adding 8 μL MT (10nM) and the absorbance of the samples was measured at a wavelength of 410 nm for 5 minutes. A negative control was added replacing BPTI with 304 μL buffer (10 mM Tris, pH8, 1 mM CaCl_2_). The range of concentrations of BPTI was adapted when the determined *K*_i_ was not in the range of these BPTI concentrations.

For MT, the *K*_i_ could be determined from **Eq. 4**, as described previously (63).

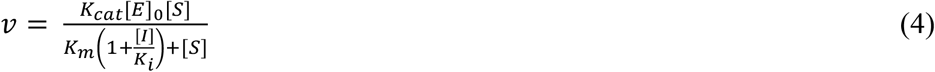

The change in experimental binding energy ΔΔ*G*_bind_ was calculated from **Eq. 5** using the *K*_i_ of the WT and the mutant, the temperature *T* at which the affinity was measured and the ideal gas constant *R*.

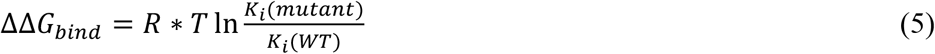

### Analysis of additivity and cooperativity

For each double mutation with available ΔΔG_bind_ prediction, we calculated the interaction energy between the two single mutations according to eq. (1).

The mutation was defined as exhibiting negative epistasis if ΔG_i_ was negative within the uncertainty of the predictions, that is:

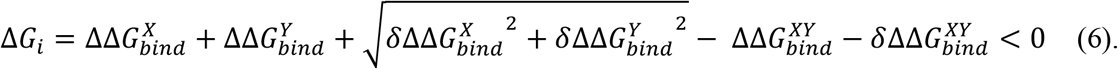

Here, 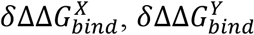 and 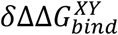 are uncertainties in prediction of ΔΔ*G*_bind_ for mutation X, Y, and XY, respectively.

The mutation was defined as exhibiting positive epistasis if ΔG_i_ was positive within the uncertainty of the predictions, that is:

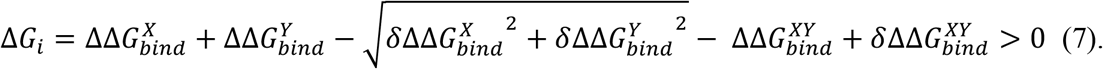

If the value of ΔG_i_ fell between the values of positive or negative epistasis the mutation was defined as additive.

