## Supplementary Information for "Climbing up and down binding landscapes: a high-throughput study of mutational effects in homologous protein-protein complexes"

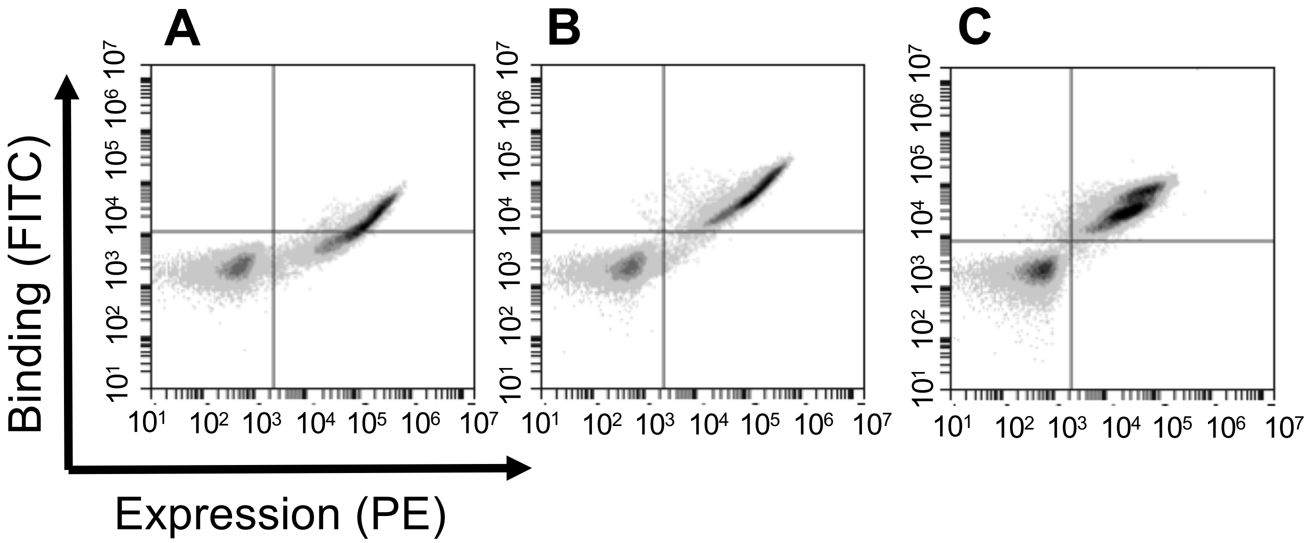


Figure S1: FACS data of $BPTI_{WT}$ binding to A) 100 nM MT, B) 20 nM ChT, C) 5 nM BT. Expression of $BPTI_{WT}$ was monitored via fluorescence of PE while binding to the three proteases was monitored via FITC fluorophore conjugated to the trypsin.

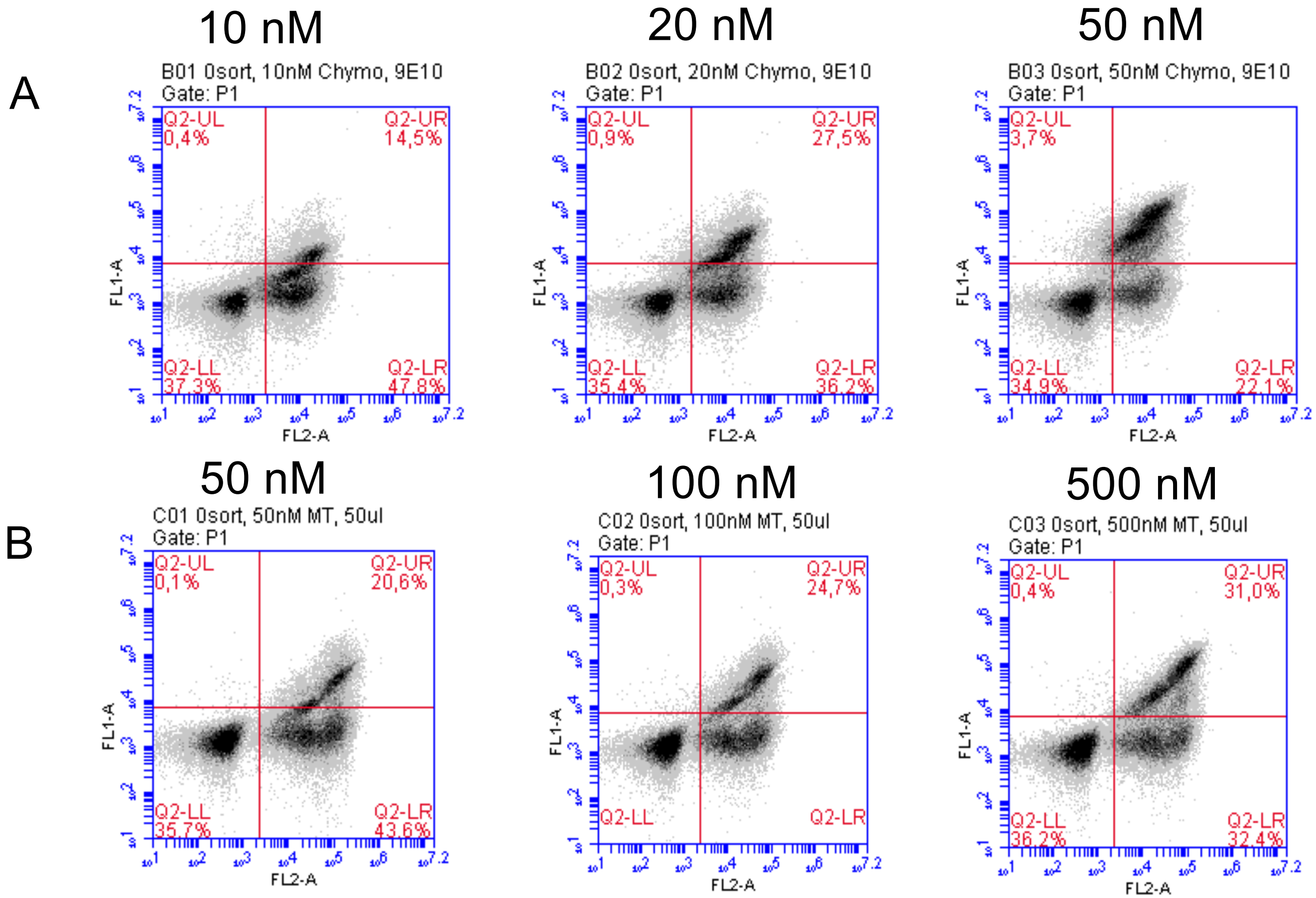


**Figure S2: Optimizing protease concentration for YSD selection.** Various concentrations of ChT (A) and MT (B) were incubated with the BPTI library and the FACS signal was measured. The concentration for BT was optimized in our previous work to be 5 nM (Heyne et al). Concentration for sorting was set to 20 nM for ChT and 100 nM for MT.

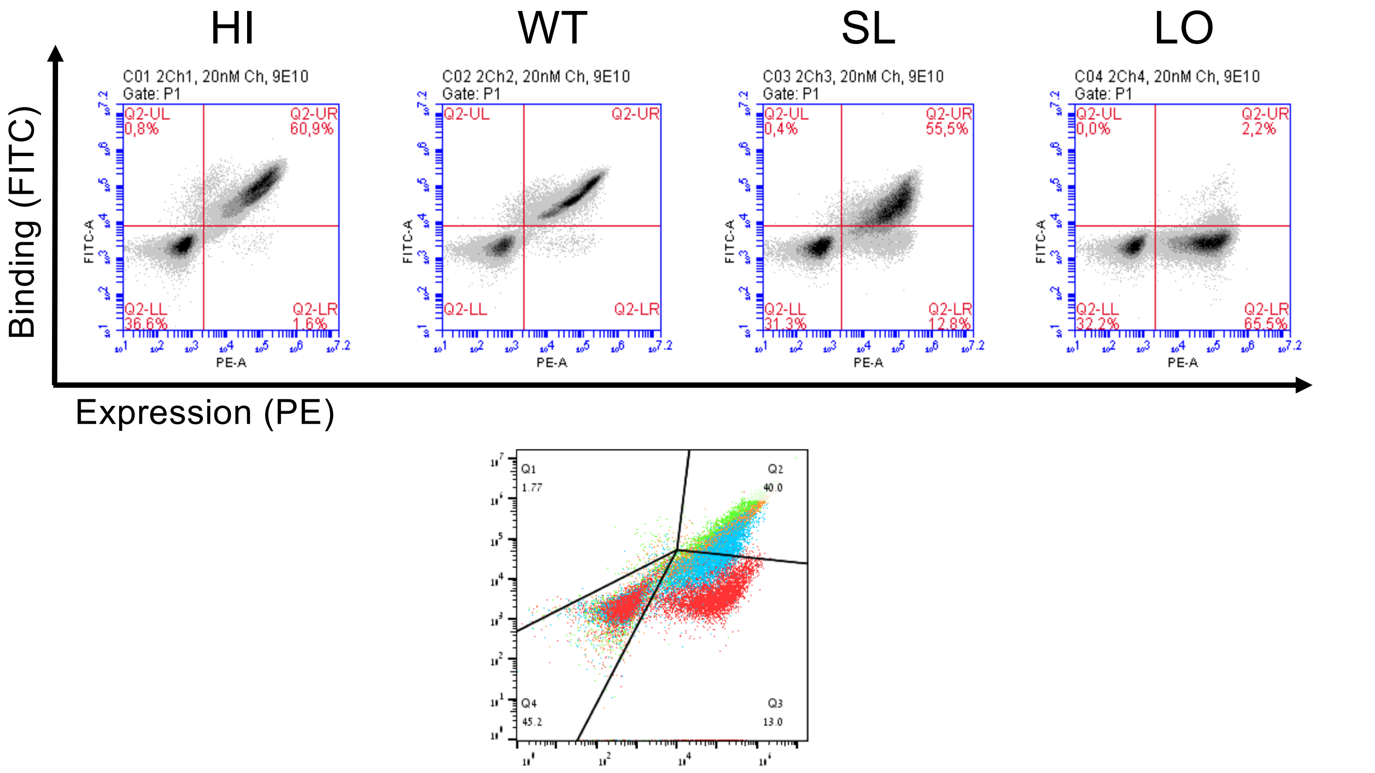


**Figure S4** Upper panel: Separation of BPTI mutant clones into four affinity windows when binding to ChT. The FACS analysis after yeast cells were sorted into 4 gates and each sorted population of cells was re-grown. Lower panel: superposition of the four populations showing cells from the LO gate in red, from SL gate in blue, from WT gate in orange and HI gate in green.

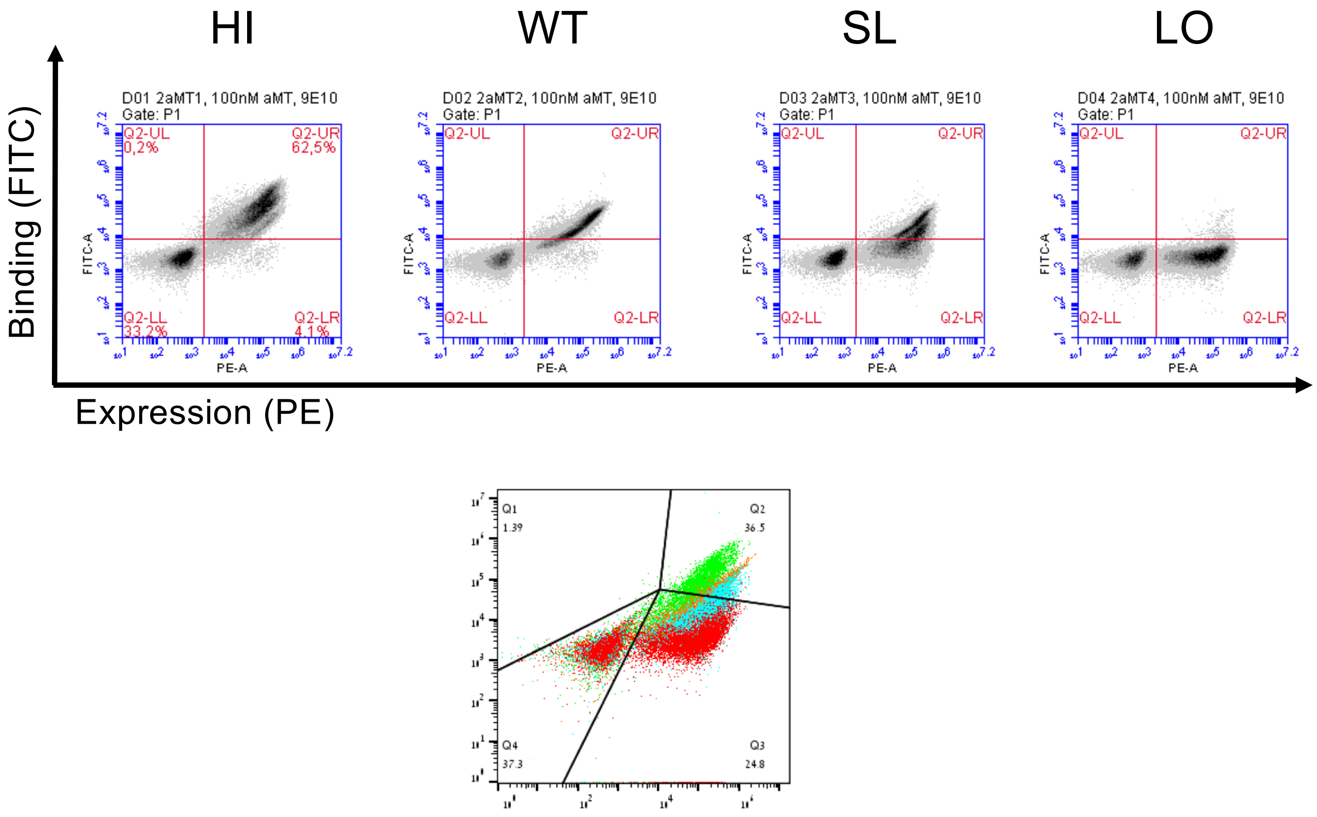


**Figure S5** Upper panel: Separation of BPTI mutant clones into four affinity windows when binding to MT. The FACS analysis after yeast cells were sorted into 4 gates and each sorted population of cells was re-grown. Lower panel: superposition of the four populations showing cells from the LO gate in red, from SL gate in blue, from WT gate in orange and HI gate in green.

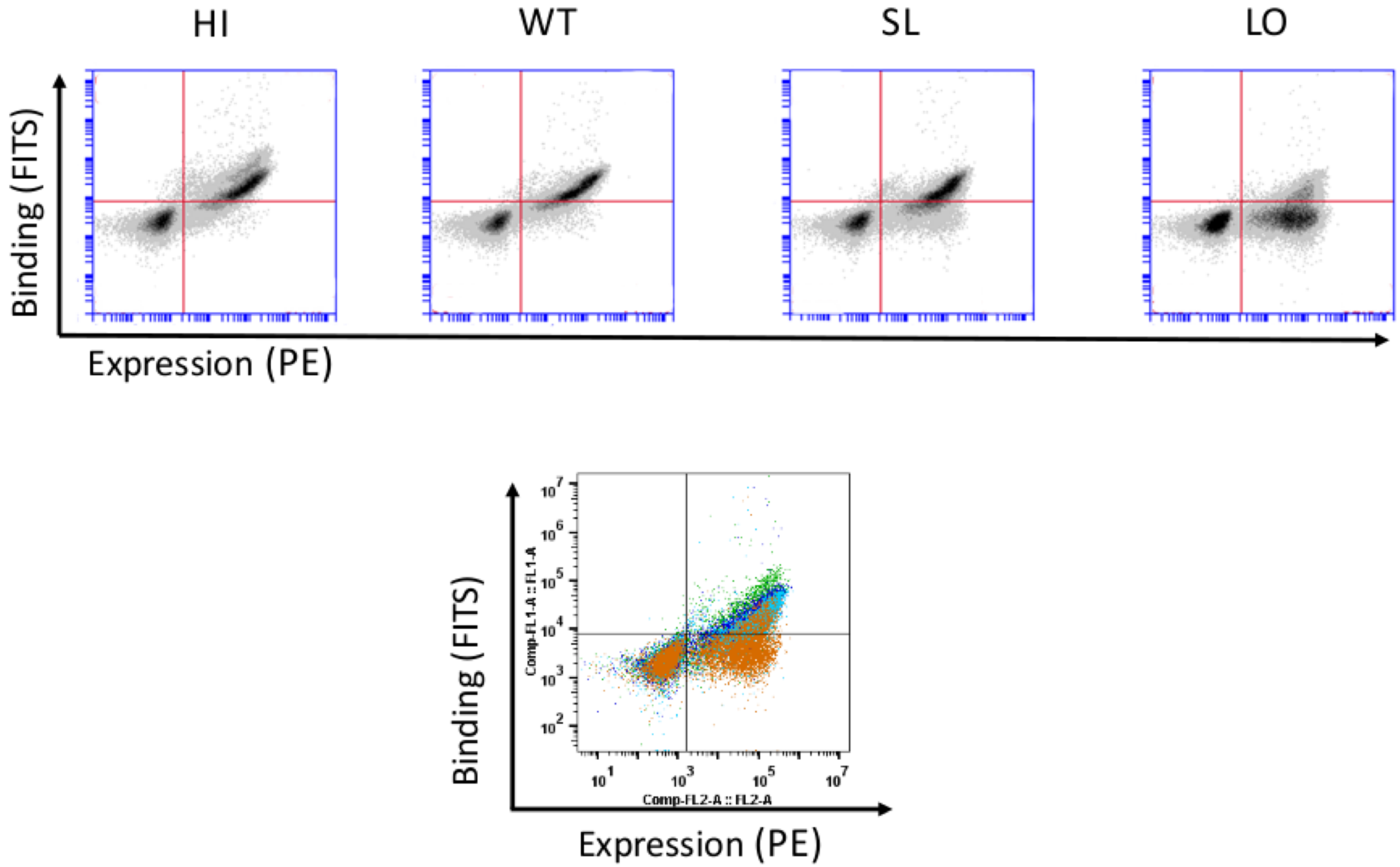


**Figure S5** Upper panel: Separation of BPTI mutant clones into four affinity windows when binding to BT. The FACS analysis after yeast cells were sorted into 4 gates and each sorted population of cells was re-grown. Lower panel: superposition of the four populations showing cells from the LO gate in orange, from SL gate in light blue, from WT gate in dark blue and HI gate in green.

(A)

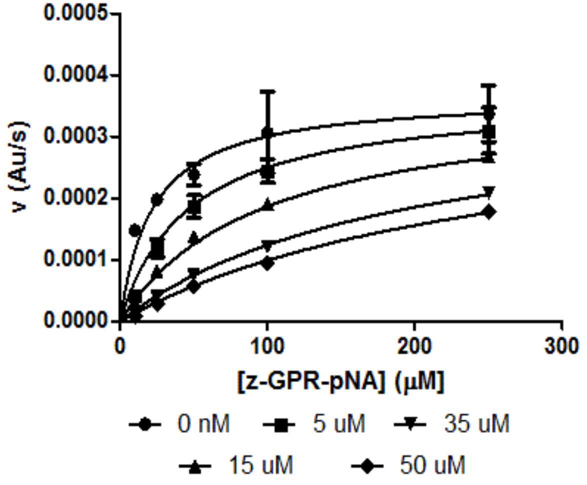


(B)

| Mutant | $K_i$ ± S. D. (µM) | $\Delta\Delta G_{exp}$ (kcal/mol) |
|---|---|---|
| WT | 4.49 ± 0.57 | 0.00 ± 0.13 |
| T11N | 32.02 ± 5.12 | 1.21 ± 0.09 |
| P13L | 6.46 ± 0.52 | 0.22 ± 0.05 |
| P13Y | 2.33 ± 0.28 | -0.40 ± 0.07 |
| K15R | 1.14 ± 0.17 | -0.81 ± 0.09 |
| R17L | 0.79 ± 0.04 | -1.07 ± 0.03 |
| I18K | 12.51 ± 1.97 | 0.63 ± 0.09 |
| V34D | 0.22 + 0.03 | -1.86 ± 0.08 |
| V34H | 6.97 + 0.68 | 0.27 ± 0.06 |
| V34W | 9.26 + 0.90 | 0.45 ± 0.06 |
| R39I | 1.50 + 0.15 | -0.68 ± 0.06 |
| T11N+V34H | 13.48 ± 2.07 | 0.68 ± 0.09 |

**Figure S6: Measurement of $\Delta\Delta G_{bind}$ values for purified MT binding to BPTI mutants**

(A) Representative experiment for determination of Ki values of BPTI mutants for MT. $BPTI_{WT}$ was used in this experiment and similar experiments were repeated with all BPTI mutants. MT cleavage of the peptide substrate Z-GPR-pNA was measured in the absence and in the presence of inhibitor, $BPTI_{WT}$. The experiment was repeated using several concentrations of $BPTI_{WT}$ as shown at the bottom. (B) Experimental results for the binding energy between MT and the BPTI variants determined in this study and used for normalization of enrichment values for MT.

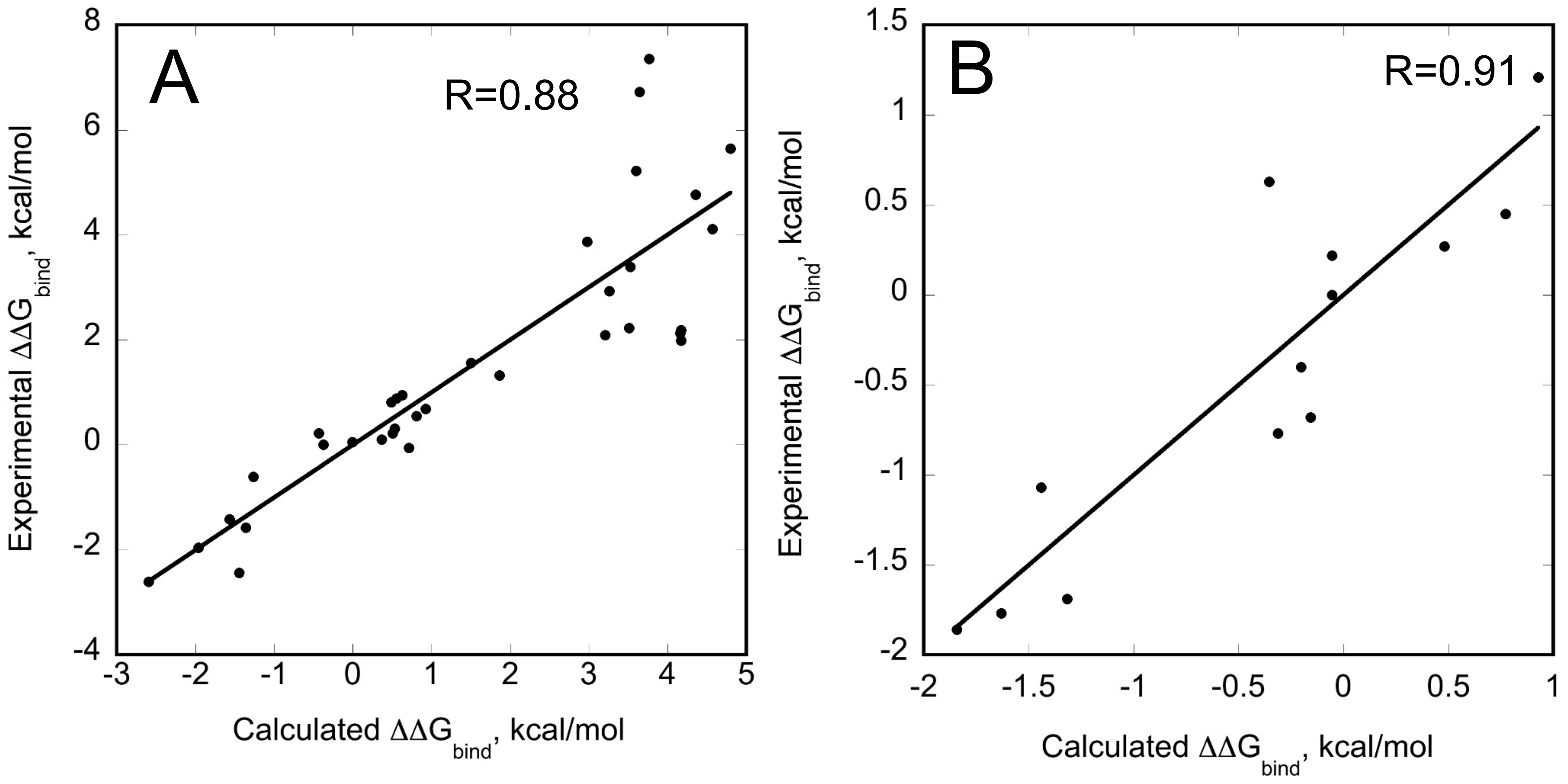


**Figure S7: Correlation between $\Delta\Delta G_{bind}$ predicted from NGS and experimental $\Delta\Delta G_{bind}$ measured on purified proteins.** (A) BPTI/ChT interactions; (B) BPTI/MT interactions. The best normalization formula was obtained that converts the enrichment value from 4 gates into the $\Delta\Delta G_{bind}$ value for each protease. The correlation coefficient R is shown on the graph. The correlation for the BPTI/BT interaction is reported in our recent paper (ref. 45).

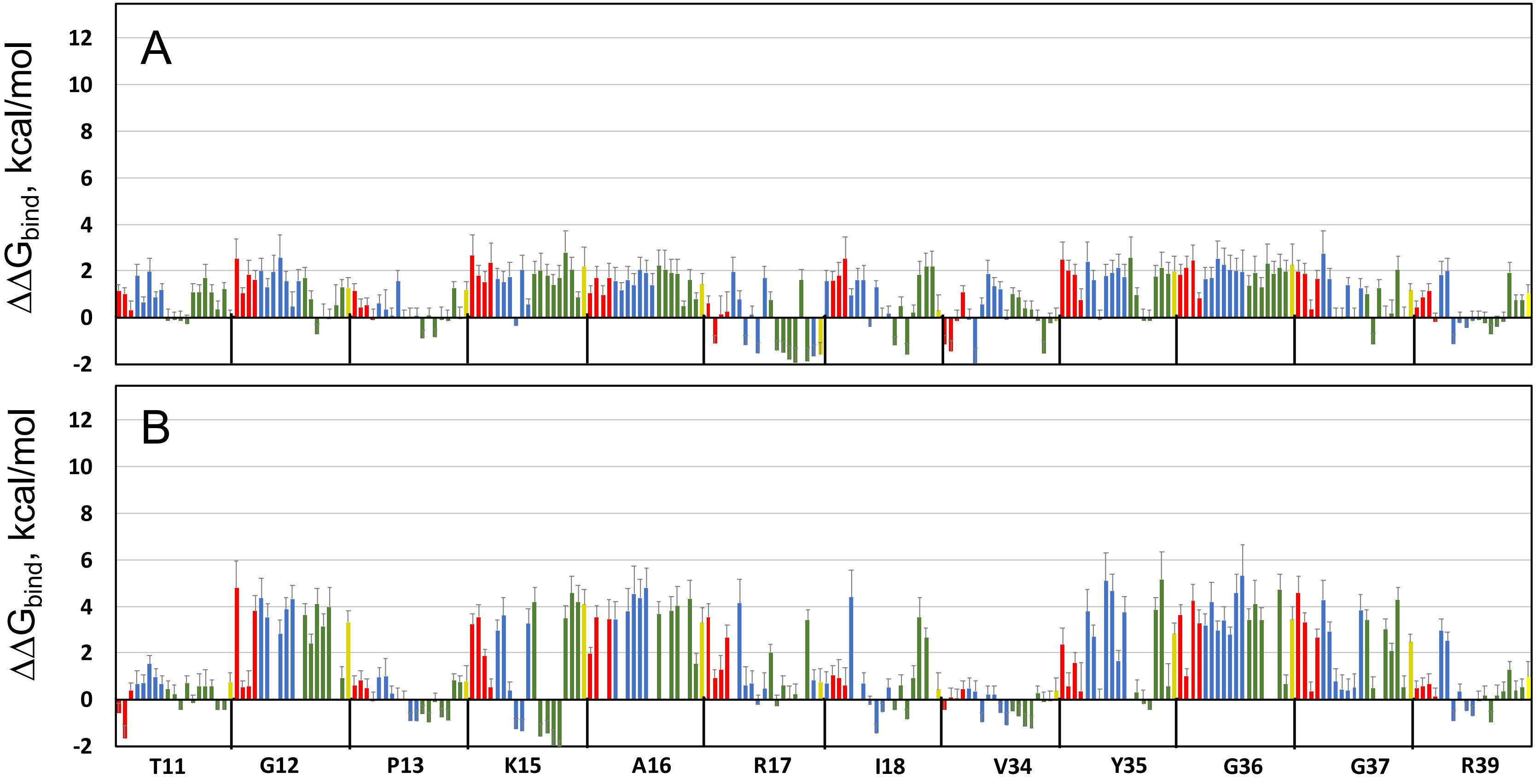


**Figure S8:** Changes in $\Delta\Delta G_{bind}$ for all single mutants of BPTI interacting with MT (A) and ChT (B). Each bar represents a mutation to one amino acid including hydrophobic amino acids (green), polar amino acids (red), charged amino acids (blue) and Cys (yellow). X-axis shows WT residue followed by position. Error bars represent the 95% CI. A similar Figure for BPTI/BT interaction can be found in our recent study (Heyne et al, ref. 45).

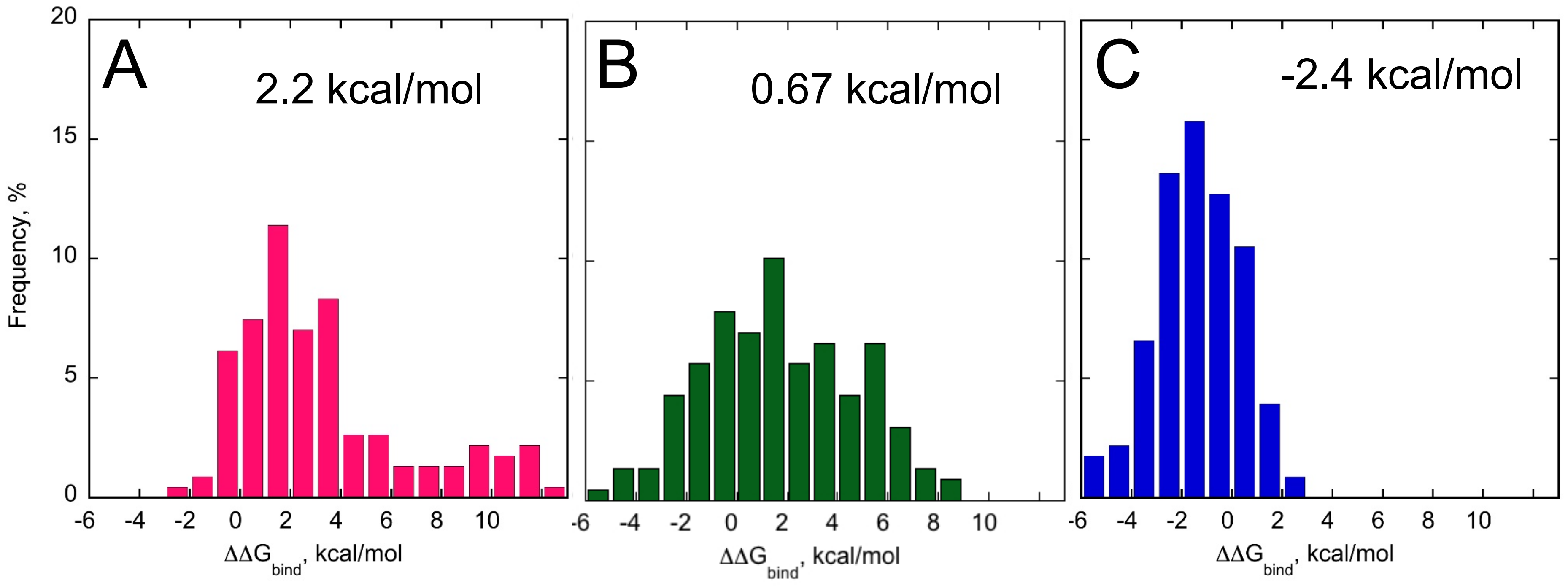


**Figure S9: Histograms of $\Delta\Delta G_{bind}$ values due to single mutational steps for BT interacting with BPTI_K15R, BPTI_A16S and BPTI_K15A** (A) BPTI_K15R mutation (B) BPTI_A16S mutation (C ) BPTI_K15A mutation. Mean value for $\Delta\Delta G_{bind}$ for each histogram is displayed on each graph. 57%, 67%, 68% of all single mutational steps are summarized on the background of K15R, A16S, and K15A BPTI mutations, respectively.

A

| 12 | 13 | 15 | 16 | 17 | 18 | 34 | 35 | 36 | 37 | 39 | |
|---|---|---|---|---|---|---|---|---|---|---|---|
| 3.9 | 3.5 | 3.7 | 3.6 | 3.2 | 3.4 | 2.8 | 2.8 | 3.6 | 2.9 | 2.3 | **11** |
| | 3.3 | 6.4 | 4.1 | 2.9 | 2.5 | 1.8 | 3.0 | 3.1 | 3.2 | 2.9 | **12** |
| | | 4.8 | 4.0 | 3.4 | 3.5 | 2.9 | 2.6 | 3.4 | 3.5 | 3.1 | **13** |
| | | | 7.5 | 4.7 | 4.6 | 3.5 | 5.7 | 6.1 | 6.1 | 4.7 | **15** |
| | | | | 4.2 | 4.2 | 3.5 | 4.6 | 4.8 | 4.6 | 3.5 | **16** |
| | | | | | 2.9 | 2.5 | 2.9 | 3.7 | 3.3 | 2.7 | **17** |
| | | | | | | 2.5 | 2.7 | 3.0 | 2.5 | 2.6 | **18** |
| | | | | | | | 2.6 | 3.4 | 2.5 | 2.6 | **34** |
| | | | | | | | | 4.2 | 3.4 | 3.6 | **35** |
| | | | | | | | | | 5.9 | 3.4 | **36** |
| | | | | | | | | | | 3.1 | **37** |

B

| 12 | 13 | 15 | 16 | 17 | 18 | 34 | 35 | 36 | 37 | 39 | |
|---|---|---|---|---|---|---|---|---|---|---|---|
| 1.3 | 1.1 | 1.0 | 1.0 | 1.1 | 1.1 | 0.6 | 1.3 | 1.1 | 1.2 | 0.9 | **11** |
| | 1.2 | 2.7 | 3.3 | 1.8 | 1.5 | 1.1 | 2.3 | 2.9 | 2.2 | 1.1 | **12** |
| | | 1.2 | 1.1 | 1.0 | 1.1 | 0.7 | 1.6 | 1.1 | 1.4 | 0.8 | **13** |
| | | | 2.7 | 1.8 | 1.4 | 0.9 | 2.0 | 2.3 | 1.8 | 0.9 | **15** |
| | | | | 1.9 | 1.6 | 1.0 | 2.2 | 3.2 | 2.5 | 1.2 | **16** |
| | | | | | 1.4 | 1.0 | 1.7 | 1.9 | 1.7 | 1.0 | **17** |
| | | | | | | 1.1 | 1.8 | 1.4 | 1.5 | 1.0 | **18** |
| | | | | | | | 1.5 | 1.2 | 1.5 | 1.0 | **34** |
| | | | | | | | | 2.7 | 2.1 | 1.5 | **35** |
| | | | | | | | | | 3.0 | 1.4 | **36** |
| | | | | | | | | | | 1.3 | **37** |

C

| 12 | 13 | 15 | 16 | 17 | 18 | 34 | 35 | 36 | 37 | 39 | |
|---|---|---|---|---|---|---|---|---|---|---|---|
| 1.4 | 0.6 | 1.5 | 1.4 | 0.6 | 0.9 | 0.7 | 1.4 | 1.8 | 0.8 | 0.6 | **11** |
| | 0.9 | 1.8 | 1.7 | 0.9 | 0.8 | 0.9 | 1.7 | 2.1 | 1.3 | 0.8 | **12** |
| | | 1.1 | 1.1 | 0.6 | 0.6 | 0.7 | 1.2 | 1.3 | 1.0 | 0.6 | **13** |
| | | | 2.1 | 0.8 | 1.4 | 0.9 | 1.9 | 2.5 | 1.6 | 1.1 | **15** |
| | | | | 0.7 | 2.0 | 0.9 | 2.5 | 2.7 | 1.7 | 1.0 | **16** |
| | | | | | 0.9 | 0.8 | 0.8 | 0.9 | 0.8 | 0.5 | **17** |
| | | | | | | 0.5 | 1.1 | 1.9 | 1.0 | 0.7 | **18** |
| | | | | | | | 1.0 | 1.2 | 1.1 | 0.6 | **34** |
| | | | | | | | | 3.0 | 2.0 | 1.2 | **35** |
| | | | | | | | | | 2.4 | 1.6 | **36** |
| | | | | | | | | | | 0.9 | **37** |

**Figure S10: The absolute values of coupling energies ΔGi averaged over a pair of BPTI positions:** (A) BT/BPTI interaction; (B) ChT/BPTI interaction; (C ) MT/BPTI interaction. On the top and on the right BPTI positions are shown. The values are color coded from large (black) to zero (white) and the average value is indicated in each square.

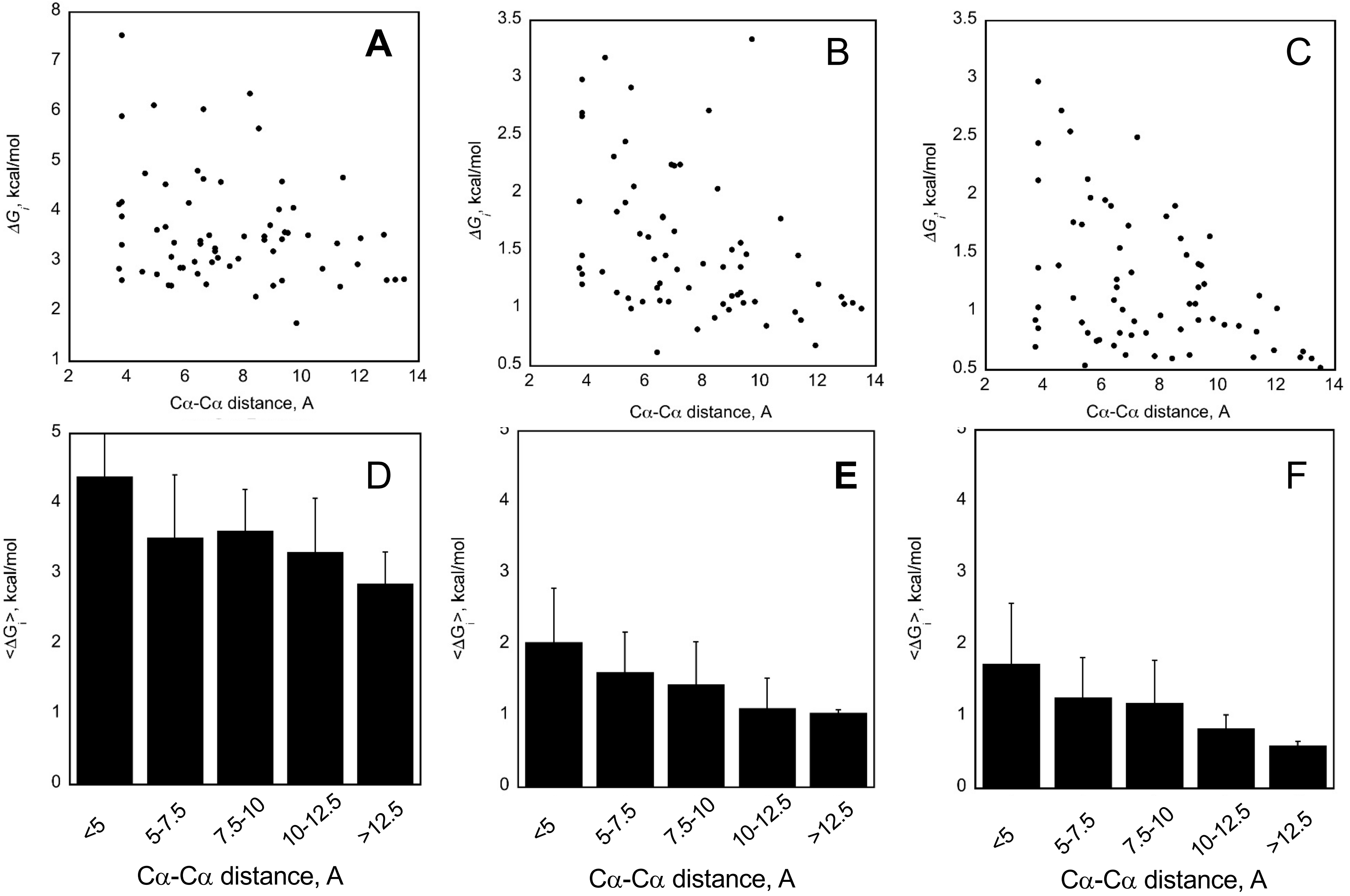


**Figure S11: Dependence of ΔGi on the Cα-Cα distance between the two positions.** Top panel shows dots that represent |ΔGi| for each pair of positions averaged over all recorded double mutations at these pair of positions. Bottom panel assigns |ΔGi| according to distance between the two Cα atoms and averages over each bin. Standard deviation for each average is shown. (A) and (D): BT/BPTI interaction; (B) and (E): ChT/BPTI interaction; ( C) and (F): MT/BPTI interaction.
